# Nanoscale indentation of plasma membrane establishes a contractile actomyosin scaffold through selective activation of the Amphiphysin-Rho1-Dia/DAAM and Rok pathway

**DOI:** 10.1101/2020.08.04.236125

**Authors:** Tushna Kapoor, Pankaj Dubey, Seema Shirolikar, Krishanu Ray

**Author notes:** Address for correspondence: Krishanu Ray, DBS, TIFR, Mumbai 400005, India., /11. Blizard Institute, Barts and The London School of Medicine and Dentistry, Queen Mary University of London, 4 Newark Institute, E1 2AT, UK.

## Abstract

Nanoscale bending of plasma membrane increases cell adhesion, induces cell-signalling, triggers F-actin assembly and endocytosis in tissue-cultured cells. The underlying mechanisms are not very well understood. Here, we show that stretching the plasma membrane of somatic cyst cell around rigid spermatid heads generates a stable, tubular endomembrane scaffold supported by contractile actomyosin. The structure resembles an actin-basket covering the bundle of spermatid heads. Genetic analysis suggests that the actomyosin organisation is nucleated exclusively by the Formins, Diaphanous and DAAM, downstream of Rho1, recruited by the Bin-Amphiphysin-Rvs (BAR)-domain protein, Amphiphysin, around the spermatid heads. Actomyosin activity at the actin-basket gathers the spermatid heads into a compact bundle and resists the invasion of the somatic cell by the intruding spermatids. These observations revealed a new response mechanism of nanoscale bending of the plasma membrane, which generates a novel cell adhesion strategy through active clamping.

**Highlights:**

- Stretching the plasma membrane around a spermatid head recruits Amphiphysin and Rho1.
- Rho1 activation triggers F-actin assembly *in situ* through Diaphanous and DAAM.
- Rho1-Rok activation assembles actomyosin scaffold around the folded plasma membrane.
- Contractile actomyosin enables plasma membrane to clamp onto the spermatid head.

**Author summary:** Sperm released from the somatic enclosure is essential for male fertility. During differentiation, the somatic cell membrane, associated with dense F-actin scaffold, tightly hold each spermatid head before release. Kapoor et al., showed that the bending and stretching of the plasma membrane trigger the assembly of an actomyosin scaffold around the bent membrane, which clamps the spermatids together preventing the premature release and somatic cell penetration. This finding provides new insight into the molecular networks activated by mechanical bending of the plasma membrane.

## Introduction

The plasma membrane is the ultimate barrier in eukaryotic cells. It also acts as a mechanical force sensor and can withstand severe distortions (Pontes et al., 2013) and indentation up to a few microns by nano-sized objects (Yang and Saif, 2007). Nanoscale deformation of the cell membrane increases cell adhesion to the substrate (Viswanathan et al., 2015), and confers cell fate and induces osteogenic differentiation of mesenchymal stem cells (Abagnale et al., 2015). It is also suggested that some viruses are likely to exploit this property to invade the host cell (Wiegand et al., 2020). Therefore, understanding the mechanism of cellular response to plasma membrane indentation is critical to identify the determinant factors.

Nanoscale bending of plasma membrane acts as hotspots for clathrin-dependent endocytosis (Zhao et al., 2017), which is likely to signal the cell and provide access to pathogens. The bending of the plasma membrane due to the tight wrapping of ∼100 nm diameter nanopillars is sensed by Clathrin, which induces dynamin-mediated endocytosis (Zhao et al., 2017). On the other hand, the bending of the basolateral cell membrane around nanopillars of diameters ∼100-400 nm triggers local F-actin assembly through the recruitment of Formin-Binding-Protein-17 (FBP-17) and Arp2/3 activation (Lou et al., 2019). Thus, molecular response to nano-indentation of the plasma membrane appears to depend on the surface topography. The underlying mechanism is, however, not investigated. Moreover, all these studies are carried out *in vitro* using synthetic conditions. Hence, these conjectures and their biological relevance are yet to be established *in vivo*.

In *Drosophila* testis, 64 spermatids mature synchronously within a somatic cell enclosure formed by the head-cyst-cell (HCC) and tail-cyst-cell (TCC) that are held together by septate junction (SJ) proteins (Dubey et al., 2019). Each mature spermatid head is a rigid, ∼10 µm long rod of ∼200 nm diameter. As the cyst matures, it moves through the terminal epithelium (TE) region at the base of the testis until mature spermatids are released from the cyst enclosure, and progress into the seminal vesicle (SV) (Dubey et al., 2016). During this last stage, spermatid heads push against the plasma membrane of the HCC and an actin-rich structure (actin cap), contributed by the somatic cyst cell, appeared to hold the spermatid heads and prevent premature release by rupturing the HCC (Desai et al., 2009; Dubey et al., 2016). Actin cap is composed of two distinct domains that are clearly separated from each other - a relatively stable actin-rich ‘basket’ domain envelopes the sperm head bundle, whereas the ‘caplets’ form around the HCC membrane whenever a sperm head intrudes beyond 2 µm from its resting position (Dubey et al., 2016). Every attempt of a spermatid head intrusion is immediately repulsed by transiently assembled caplets which are formed by the WASp-Arp2/3-dependent process around the indented plasma membrane of the HCC (Dubey et al., 2016). The caplet disappears as soon as the intruding spermatid head reverts to its initial position. The mechanism of the basket assembly, which persists even after the spermatid release, was not explored.

Here, we provide a comprehensive sub-cellular and molecular-genetic evaluation of the basket assembly that helped to derive a novel clamping mechanism. We find that the basket is composed of a highly tubular membrane scaffold supported by actomyosin. It is assembled exclusively through concerted activities of the Rho1 GTPase, and the formins Diaphanous and DAAM. The Bin-Amphiphysin-Rvs (BAR) domain protein, Amphiphysin, possibly recruits Rho1/RhoA at the basket. Rho1 recruits and activates nonmuscle myosin-II through Rho-kinase. The actomyosin activity maintains the integrity of the actin-basket and holds the spermatid heads together in a parallel bundle, which is essential for maintaining male fertility (Dubey et al., 2019). Basket domain also prevents the penetration of the somatic cell by spermatid heads. Together, these results describe an unprecedented role of membrane bending in triggering the assembly of an actomyosin clamp at the plasma membrane. We postulate that the bending of the HCC plasma membrane around spermatid heads triggers the recruitment of specific BAR domain proteins to initiate the basket assembly, and discuss the significance of these findings in the context of wound healing and tissue morphogenesis.

## Results

### Actin-Basket consists of extensively tubular endomembrane and F-actin

The cysts containing coiled spermatids, with characteristic compacted NBs (Fig 1A; arrows, Fig 1B) are found within the TE zone. During the initial stage of individualisation, F-actin cones (Investment Cones – IC) consisting of dendritic actin form around each spermatid head (arrowhead, Fig 1B; Fig S1A), and move down the tail (Noguchi and Miller, 2003). As the ICs progressed, the spermatid heads were rearranged in a parallel bundle (Nuclei Bundle – NB, Fig S1B-E), and a different type of F-actin structure, called actin cap (Desai et al., 2009), appears inside the HCC around the NB (yellow arrowheads, Fig S1C-E). Actin cap consists of a dense F-actin basket and multiple caplet domains (Dubey et al., 2016) (arrowhead and arrow, Fig S1F). The basket is most pronounced in the middle region of the TE, which had fully coiled spermatid tails (arrowhead, Fig S1F).

**Figure 1.**
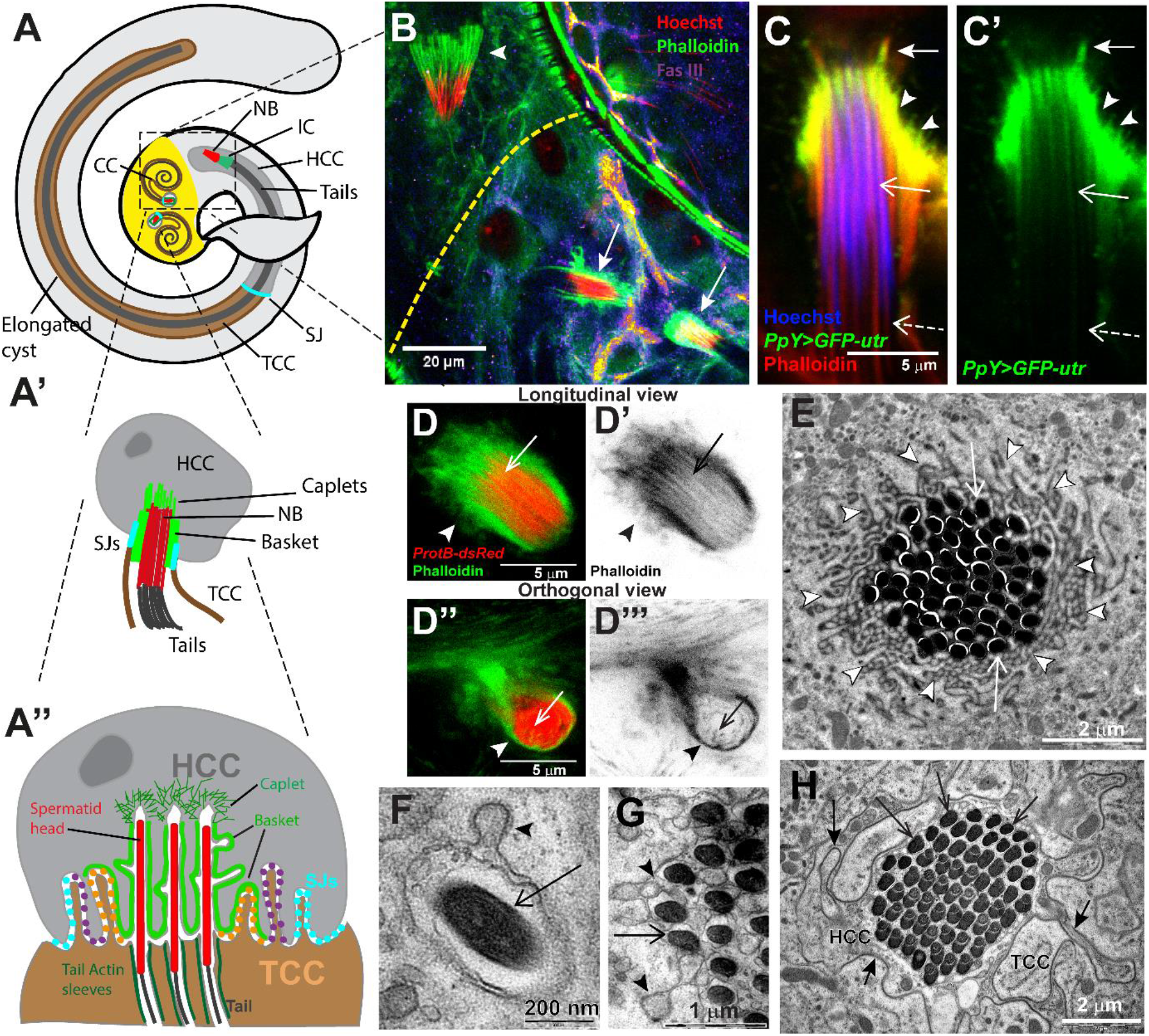
Network of tubular, actin-rich membrane covers individual spermatid heads within a cyst during the final stages of sperm maturation in *Drosophila* testis. **A-A’’)** Schematic of *Drosophila* testis. Sixty-four spermatids synchronously develop within the cyst enclosure. After individualisation, the elongated spermatid tails (dark grey) coil up inside the cyst and move to the TE region (yellow zone) at the base of the testis. The nuclei bundles or NBs (red) of coiled spermatids are covered by an actin cap (green), consisting of basket and caplet domains. **A’** and **A”** depicts the expanded views of the arrangement inside the HCC. As depicted in **A”**, each spermatid head is enclosed with an actin-rich membrane sleeve. The HCC-TCC interface is highly folded, and marked by the SJs. **B)** Confocal image of a wild-type testis, stained with Hoechst dye (red), marking the spermatid heads, and Phalloidin (green) marking F-actin. Actin-caps form around the spermatid head bundles (NB, red) (arrows) once the cyst is inside the TE region. The yellow dashed line marks the boundary of TE region marked by the Fas-III immunostaining (Fire LUT). Arrowhead indicates the spermatid head bundle of an early individualising cyst, with associated IC at the caudal end of the NB outside the TE region. **C-C’)** *PpY>GFP-utr* (green) expression in cyst cells, stained with Hoechst (blue) and Phalloidin (red). The caplets (arrow) and the basket domains (arrowheads), as well as a part the actin sleeves (open arrows), are marked by the GFP-Utr. **D-D’’’)** STED microscopy images of NB (red) and associated actin cap (green) in *ProtB-dsRed* expressing testis, stained with Phalloidin. The longitudinal view (D-D’) shows that the actin structure is present in between the packing of individual sperm heads (arrows). Orthogonal section (D’’-D’’’) through the NB also depicts the presence of actin in the region between individual sperm heads (arrows). Arrowheads mark the basket actin surrounding the entire bundle. **E-H)** TEM images of transverse sections of NBs from wild-type testes depict spermatid heads (open arrows, E) surrounded by extensively folded and tubular membrane (arrowheads, E) near the tips of the spermatid heads, which are buried relatively deeper inside the HCC. Such tubular membrane was not visible at the level of the HCC-TCC junction (arrows, F). High magnification images of individual spermatid heads inside the HCC (G) shows two layers of plasma membrane surrounding each spermatid head. The outer layer, likely to be contributed by the HCC, have tubulobulbar extensions (arrowheads, H).

To get a more comprehensive understanding of the actin cap, which is formed inside the HCC (Desai et al., 2009), we used a combination of confocal and Structured Emission Depletion (STED) microscopy. We used *PpYGal4*, which expresses at a relatively higher level in the HCC during the terminal stage of spermatid differentiation. The *PpY>GFP-utr* expression marked caplets (arrows, Fig 1C-C’), as well as a dense F-actin basket, surrounding the NB (arrowheads, Fig 1C-C’). In addition, we noted F-actin sleeves around individual spermatids (open arrows, Fig 1C-C’), henceforth referred to as ‘actin sleeves’. The actin sleeves present in between each spermatid head in an NB were also prominent in the STED images (open arrows, Fig 1D-D’’’). The caudal part of the actin sleeves, marked by Phalloidin, but not by GFP-Utr (dashed open arrow, Fig 1C-C’; open arrows, Fig 1D’’-D’’’) are most likely contributed by the TCC, since mature sperms released into the SV do not stain for F-actin.

As described earlier (Tokuyasu et al., 1972b), Transmission Electron Microscopy (TEM) of transverse sections of NBs indicated that individual spermatid heads are tightly wrapped by the HCC membrane (arrows, Fig 1E). The HCC membrane is highly tubulated at the rostral end of the spermatid heads (arrowheads, Fig 1E), which are buried deep into the HCC. The membrane tubules also had bulbous termini (arrowheads, Fig 1F-G), and electron-dense material coated the membrane buds (arrowhead, Fig 1F). These structural features resembled the tubulobulbar-complex found around spermatid heads inside Sertoli cells in mammalian testis (Guttman et al., 2004). Such tubules were not detected near the HCC-TCC interface (arrows, Fig 1H), which is held together by SJs (Dubey et al., 2019). Together, these analyses indicated that each spermatid is tightly sleeved with the somatic cell membrane supported by F-actin (actin sleeve), and the head bundle (NB) is surrounded by a highly tubular actin-rich membrane, which we redefine as the ‘basket’ domain (Fig 1A”) (Video S1).

The baskets were classified in two categories - *mature* baskets with distinctively rich Phalloidin staining (Fig 1C; S1F), and relatively *thin* ones with barely detectable Phalloidin staining surrounding the NB (Fig S1G-G’). These two morphological classes were also distinguished by significantly different F-actin volume (Fig S1H). Altogether, these observations indicated that the basket domain forms around the spermatids heads as the coiled spermatids progress into the TE.

### Basket domain is a contractile structure composed of relatively stable F-actin

*Ex-vivo* time-lapse imaging of *Drosophila* testes expressing *sqh::utr-GFP* and *ProtB-dsRed*, marking the F-actin and the NB, respectively, revealed that the basket periodically contracts and expands along the axis parallel to the NB (Fig 2A; Video S2). To understand the effect of periodic basket contraction, we used high-intensity pulsed IR laser (800 nm) to ablate a part of the basket. Focussing the laser beam in a 0.75 µm^2^ (10×10 pixels) area near the edge of the basket region caused an isotropic retraction of F-actin from the centre of the spot (asterisk, Fig 2B, B”). Over time, the F-actin surrounding the NB dispersed (follow arrowheads, Fig 2B) and accumulated at the caplet region (arrow, 30’ 56”, Fig 2B; Video S3), with concomitant sheering of the NB (Fig 2B’). To ensure that the loss of F-actin from the basket region caused NB disruptions, we ablated a control spot in the HCC cytoplasm, away from the basket (dashed region, Fig 2C-C’) using the same laser power. It did not perturb the caplet and the basket (arrows and arrowheads, Fig 2C), as well as the NB organisation (Fig 2C’; Video S4). These results suggested that that basket is a contractile structure required for maintaining the spermatid heads in a tight bundle.

**Figure 2.**
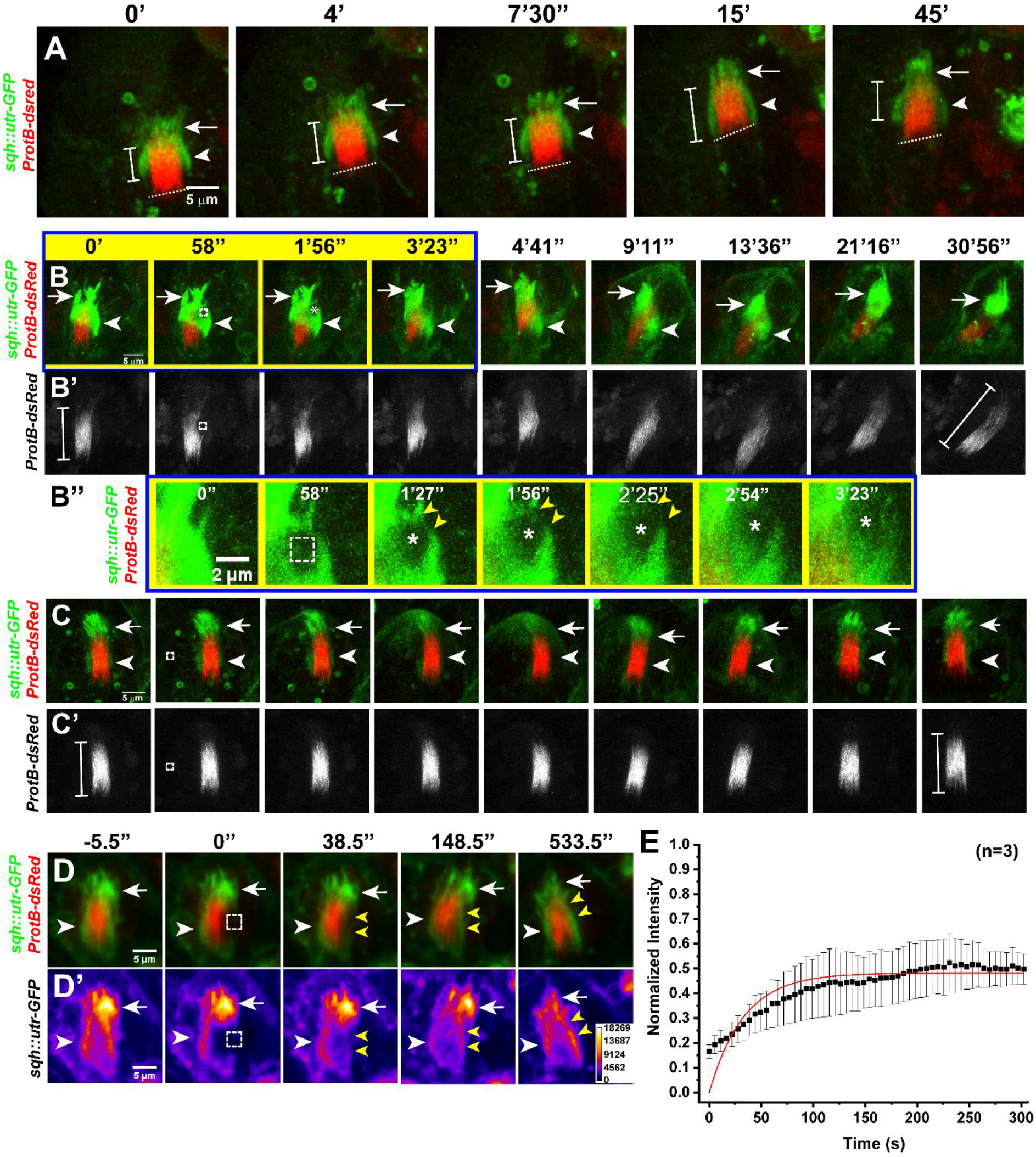
Basket domain maintains the spermatid heads in a bundle. **A)** Time-lapse images of *sqh::utr-GFP; ProtB-dsRed* expressing testis, depicting the actin cap (green) around the NB (red). Arrow indicates the caplet; the arrowheads indicate the basket domain, and the dashed line denotes the caudal end of the NB. Note that the length of the basket, indicated by the scale bar, oscillates along the long axis. **B-C)** Time-lapse images after part ablation (dashed outline) of the basket domain (B-B’’), and a control spot in the HCC cytoplasm (C-C’), of *sqh::utr-GFP; ProtB-dsRed* testes, using 800 nm laser. Arrows mark the caplets; the arrowheads mark basket and asterisks mark the centre of the ablated ROI. Closed lines indicate the length of the NB, before ablations, and at 30’56”. (B”) High magnification images of 0-3’23” frames of B, showing the expanding edges of the ablated region (yellow arrowheads). **D)** Fluorescence recovery after photobleaching (FRAP) of *sqh::utr-GFP* in the basket domain. The intensity profile is depicted in a false colour heat map (FIRE, D’). The dashed outline marks the photobleached ROI; yellow arrowheads mark the photobleached zone during recovery; the arrowheads mark the basket domain, and the arrows mark the caplets. **E)** The FRAP recovery profile (mean ± S.D.) fitted with the Levenberg-Marquardt iteration algorithm.

Further, the Fluorescence Recovery After Photobleaching (FRAP) revealed that the F-actin at the basket domain is composed of stable and dynamic components (Video S5). The post bleach recovery (t= 5.5’’ onwards, Fig 2D-D’) reached saturation within 100 seconds (Fig 2E). The data were fitted to a single exponential using the Levenberg-Marquardt iteration algorithm, and the maximum recovery was found to be 0.49 ± 0.006, indicating nearly equal amounts of mobile and immobile fractions of F-actin. Hence, nearly 50% of the F-actin at the basket domain appeared to remain stable for a considerable period.

We also noted that thin F-actin-rich extensions (actin tubules) occasionally emanate from the basket domain (open arrowheads, Fig S2; Video S6). These structures may contribute to the partly dynamic nature of the basket, and may potentially mark the tubular membrane extensions observed by the TEM (Fig 1E, G).

Together, the anatomical data and the time-lapse analyses suggested that a relatively stable membrane-actin assembly around the spermatid heads during the final stages of spermatid maturation maintains them in a tight bundle. We wanted to understand how such a complex and unprecedented structure forms and its biological relevance.

### Rho1 activation organises the F-actin assembly at the Basket domain

*Drosophila* genome codes for Rho1 (Rho A in mammals), three different Rac isoforms, and a Cdc42 orthologue (www.flybase.org). To understand the molecular mechanism of F-actin assembly in the basket domain, we knocked down, as well as altered the functions of these three classes of Rho GTPases in the cyst cells, using *PpYGal4*. Three quantitative parameters - the number of intact NBs, number of progressed ICs and basket morphology - were used to ascertain the cellular and biological impact of these perturbations. Loss of actin cap disrupts the NB and leads to premature release inside the testis (Desai et al., 2009; Dubey et al., 2016). Therefore, we counted the number of intact NBs at the base of the testes to determine the effects of the tissue-specific perturbations (arrows, Fig 3A). Loss of F-actin assembly during the individualisation stage, preceding the spermatid coiling, affects the IC assembly and progression, and the NB packing (Ghosh-Roy et al., 2005). Therefore, we estimated the number of progressed ICs, to clarify that the effect was restricted to the final stage of sperm maturation. Third, we analysed the basket morphology and volume marked by the Phalloidin staining, as well as the anisotropy of NB packing to understand the effects at the subcellular level.

**Figure 3.**
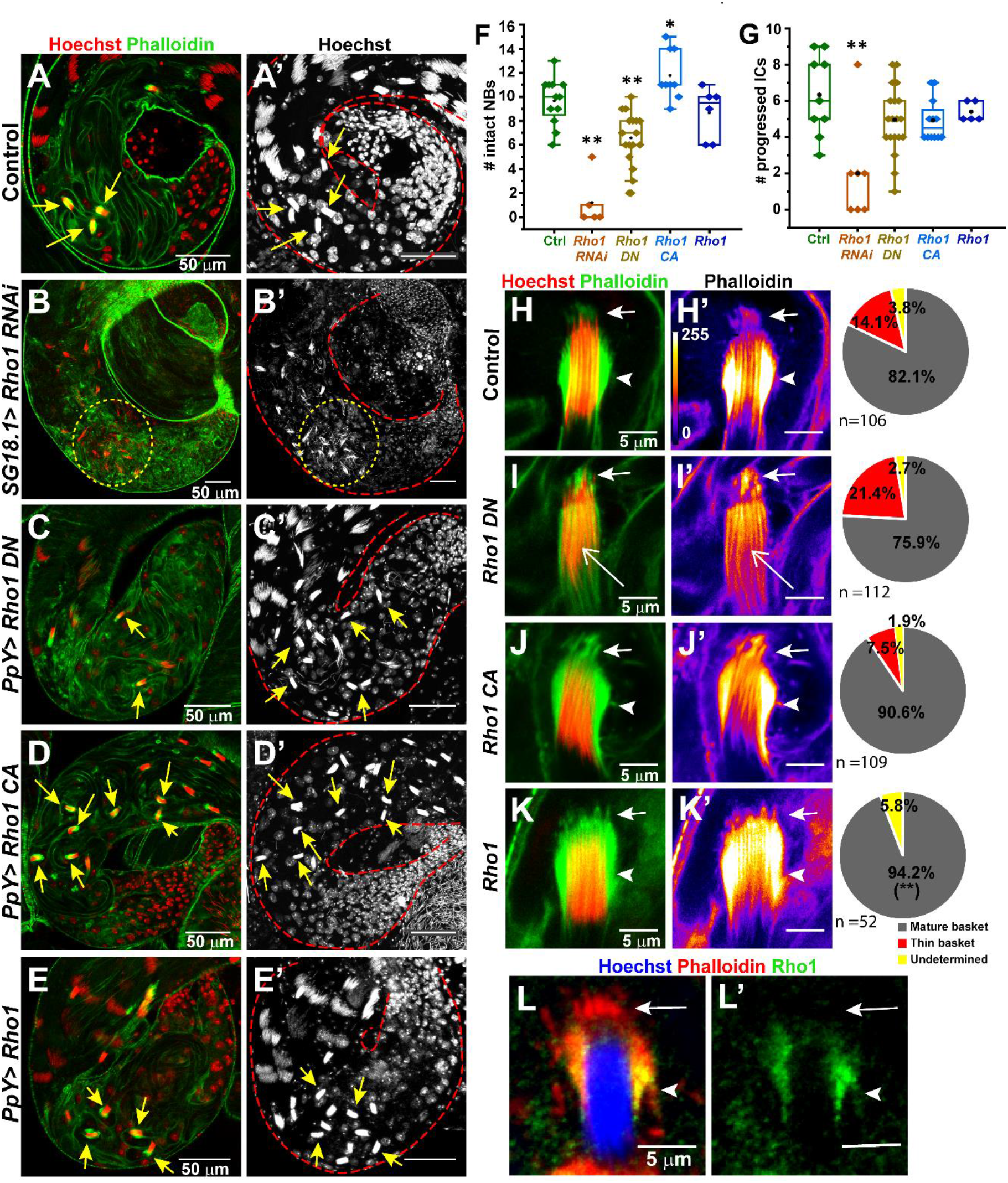
Rho 1 perturbation in the cyst cells affects the stability of the basket domain and NB. **A-E)** Confocal sections of testes base from the Control (A-A’), *SG18*.*1>Rho1* RNAi (B-B’), *PpY>Rho1N19DN* (C-C’), *PpY>Rho1V14CA* (D-D’) and *PpY>Rho1* (E-E’) backgrounds, stained with the Hoechst (red/ white) and Phalloidin (green). Arrows indicate intact NBs associated with baskets. A’-E’ represent maximum intensity projections of Hoechst channel, depicting all the NBs in the testes. **F-G)** Box plots depict the number of intact NBs (F) and the progressed ICs (G) upon Rho 1 perturbations in the cyst cells. The pairwise significance with respect to the control value was estimated using One-way Anova® and Mann-Whitney U tests, respectively, and the p-values-* <0.05, ** <0.01, *** < 0.001 are indicated on each box. **H-K**) High magnification images of an NB associated with the actin cap from the control (G-G’), *Rho 1DN* (I-I’), *Rho 1CA* (J-J’), and *Rho 1* (J-J’) expressing testes, stained with Hoechst (red) and Phalloidin (green/ FIRE). Arrows, arrowheads and open arrows indicate caplets, basket, and actin sleeves, respectively. Pie charts indicate relative proportions of different basket morphologies classified as mature (as shown in Fig S1F), thin (as shown in S1G), and undetermined. The pairwise significance with respect to the control value was estimated using Fisher’s Exact test, and the p-values-* <0.05, ** <0.01, *** < 0.001 are indicated on the pie chart. **L-L)** NB and the associated basket domain stained with anti-Rho1 (green), Hoechst (blue), and Phalloidin (red). Note that Rho1 is enriched at the basket domain (arrowhead), and not in the caplet (arrow).

The *PpY>Rho1 RNAi* combination was lethal. Knockdown of Rho1 using the relatively milder *SG18*.*1Gal4*, survived to the adult stage. The TE region contained mostly disrupted NBs (yellow dashed circle, Fig 3B-B’, F) and very few progressed ICs (Fig 3G; arrowheads, Fig S3B). The overexpression of dominant-negative Rho1-N19 using *PpYGal4* reduced the number of intact NBs (Fig 3C-C’, F) without affecting the progressed ICs (Fig 3G; arrowheads, Fig S3C). A higher proportion of these NBs was associated with visibly thin or deformed baskets (open arrows, Fig 3I-I’; pie chart), as compared to control (Fig 3H-H’; pie chart). The expression of constitutively-active Rho1-V14 (*PpY>Rho1CA*) increased the number of NBs (Fig 3D-D’, F) and the proportion of mature baskets (Fig 3J-J’; pie chart), without altering the number of progressed ICs (Fig 3G; arrowheads, S3D). Similarly, Rho1 overexpression (*PpY>UAS Rho1*) significantly increased the fraction of NBs associated with mature baskets (pie chart, Fig 3K), without altering the number of NB (Fig 3E-E’, F) and progressed ICs (Fig 3G; arrowheads, Fig S3E). Thus, Rho1 activity is implicated in the F-actin assembly at the basket domain. Finally, Rho1 immunostaining revealed a visible enrichment only at the basket domain, and not at the caplets (arrowhead and arrow, Fig 3L-L’). Together these results indicated that Rho1 localisation and activation at the basket domain are essential for the F-actin assembly.

Next, we tested the roles of Cdc42 and Rac1. The *PpY>Cdc42 RNAi* testes were shrunken, with no identifiable mature cysts (Fig S4C-D), suggesting that Cdc42 has a broader role in sperm differentiation. Overexpression of the dominant-negative *Cdc42-L89* significantly increased the average number of intact NBs (Fig S4E, P), but there were no identifiable defects in the basket morphology (Fig S4F-F’). Hence, we ruled out a direct involvement of Cdc42 in the basket assembly. The *Rac1* and *Rac2 RNAi*, as well as the overexpression of the dominant-negative Rac1-N17 and Rac1-L89, respectively, did not change the NB and IC counts (Fig S2). However, we noticed an unusually large number of actin tubules emanating from the basket in the *Rac1 RNAi* and DN backgrounds (yellow arrowheads, Fig S2H-H’, L-L’). Also, the Rac1-GFP, expressed from its endogenous promoter, was moderately enriched in and around the basket (arrowhead, Fig S4O-O’). Thus, we concluded that Rac1 is likely to maintain actin tubule dynamics. Together, these results suggested that though Rac1 is not directly involved in the basket assembly, it may fine-tune actin-membrane dynamics in the basket.

In summary, tissue-specific perturbation of different class of Rho-GTPases in the cyst cells suggested that the basket domain is exclusively formed due to the activation and recruitment of Rho1. These observations also raised two specific questions – whether the recruitment and activation of Rho1 lead to the F-actin assembly at the basket domain and how Rho1 is recruited.

### Basket is composed of linear F-actin nucleated by the formins - Diaphanous and DAAM

The association of GTP-bound Rho1 activates a sub-group of the formin class of the F-actin nucleation promotion factors (Kuhn and Geyer, 2014). Therefore, to identify the role of formins in the basket assembly, we separately expressed RNAi constructs for all the six formins coded by *Drosophila* genome (Breitsprecher and Goode, 2013), *viz*., Diaphanous (Dia), Dishevelled Associated Activator of Morphogenesis (DAAM), Formin-like (Frl), Formin 3 (Form 3), Formin homology-2 domain-containing (Fhos) and Cappuccino (Capu), using *PpYGal4*. The *dia RNAi* expression led to a severe loss of NBs (Fig 4B-B’, D) with no significant change in the number of progressed ICs (Fig 4E; S5B). We also noticed a large number of free spermatid heads in the TE in *dia RNAi* testes (yellow dashed circle, Fig 4B-B’), suggesting that NBs were disrupted after individualisation. Amongst the few remaining intact NBs, more than 35% (p = 0.0086) were associated with severely deformed and thin baskets (Fig 4G-G’; pie chart) with significantly lower volume (Fig 4I). Furthermore, the NB anisotropy estimate revealed a lower trend (i.e., less ordered arrangement), although the difference was not statistically significant (Fig 4J). The *DAAM RNAi* also caused a significant, but comparatively smaller reduction of NBs (Fig 4C-D) than *dia RNAi*, with no alteration in the number of progressed ICs (Fig 4E, S5C). Similar to the *dia RNAi*, nearly one-third of the baskets in the *DAAM RNAi* testes were thin and deformed (Fig 4H-H’; pie chart) with marginally decreased volume (Fig 4I), suggesting that DAAM may also be involved in the basket assembly. However, we did not observe a change in NB anisotropy (Fig 4J). Together, these observations implicated both Dia and DAAM in the basket assembly.

**Figure 4.**
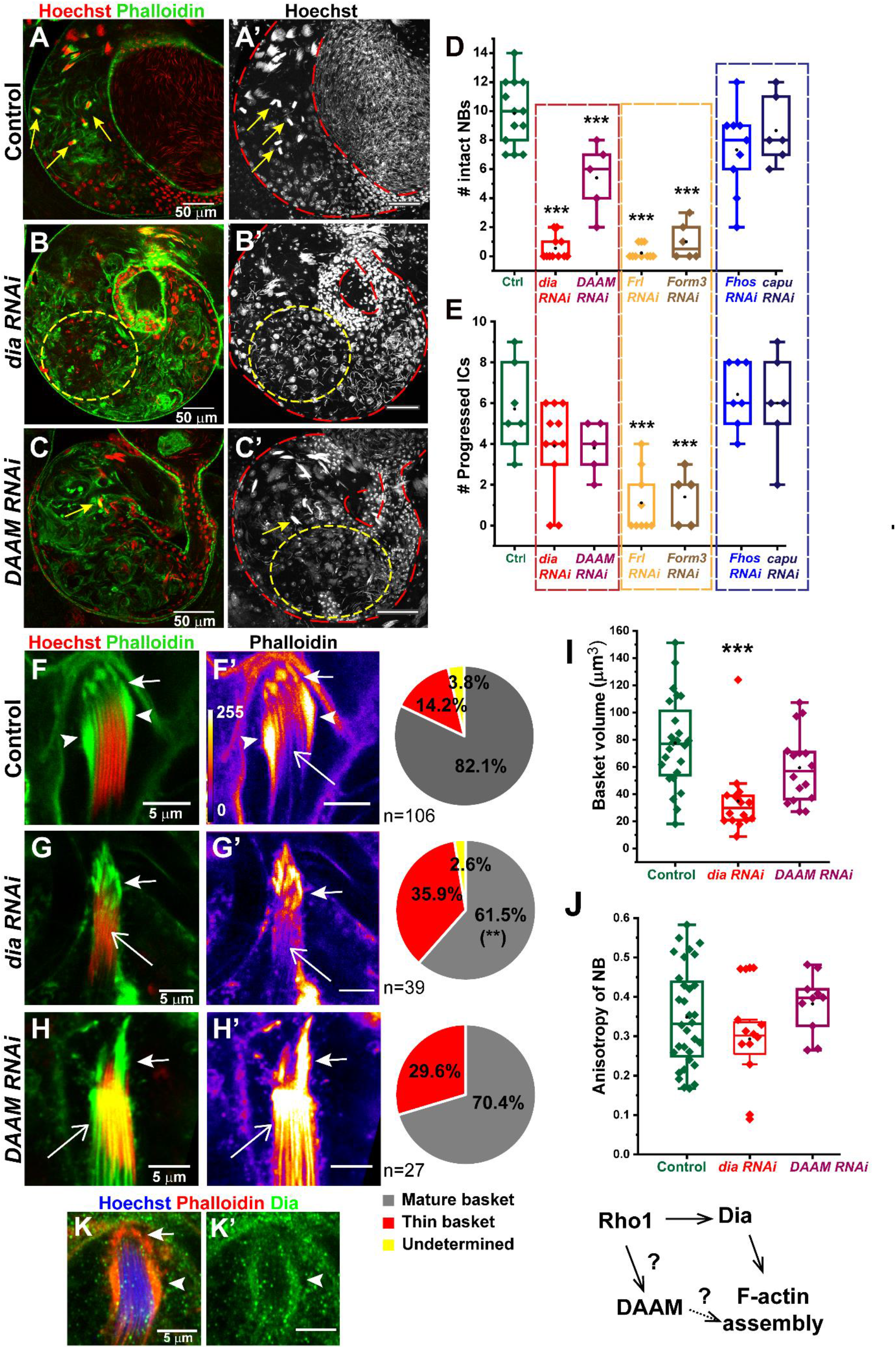
F-actin, nucleated by the formin, Diaphanous and DAAM, constitutes the basket domain. **A-C)** TE regions of *PpY>GFP* (Control, A-A’), *PpY> dia RNAi* (B-B’) and *PpY> DAAM RNAi* (C-C’) testes stained with Hoechst (red/ white) and Phalloidin (green). A’-C’ represent maximum intensity projections, depicting all the NBs in the TE region. **D-E)** Box plots depict the number of intact NBs (D) and the progressed ICs (E) on perturbation of six different formins in the cyst cells. The pairwise significance with respect to the control value was estimated using Mann-Whitney U tests, and the p-values-* <0.05, ** <0.01, *** < 0.001 are indicated on each box. **F-H)** High magnification images of NBs stained with Hoechst (red) and Phalloidin (green/FIRE), from the control (F-F’), *dia* (G-G’) and *DAAM* (H-H’) *RNAi* testes. Pie charts indicate relative proportions of different basket phenotypes. Arrows, arrowheads and open arrows indicate caplets, basket, and actin sleeves, respectively. Significance was calculated using Fisher’s exact test, and the p values - * <0.05, ** <0.01, *** < 0.001 are indicated on the pie charts. **I-J)** Box and dot plots depict the volume of the basket domain (I) and the organisation (Anisotropy) of spermatid head alignment in the NBs (J) in Control, *dia* and *DAAM* RNAi backgrounds. The pairwise significance of difference with respect to the control values was estimated using Mann-Whitney U tests and One-way Anova®, respectively, and the p-values-* <0.05, ** <0.01, *** < 0.001 are indicated on each box. **K)** NB and the associated basket domain stained with anti-Dia (green), Hoechst dye (blue), and Phalloidin (red). Note that Diaphanous is enriched at the basket domain (arrowheads), and not in the caplet (arrow).

Although *Frl* and *Form3 RNAi*s also severely reduced the number of intact NBs (Fig 4D; S5D, G), the number of progressed ICs were also reduced significantly (Fig 4E; S5F, I). A few remaining NBs that we found in these backgrounds appeared to have intact baskets (arrowheads, Fig S5E, H), but we could not quantify their morphology, as the number of intact NBs was insufficient. Hence, we concluded that the NBs were lost due to considerable disruption of an earlier differentiation process. Lastly, the expression of *Fhos* and *Capu RNAi*s did not affect NB (Fig 4D; S5J, M) and IC (Fig 4E; S5 L, O) counts. The basket morphologies were also similar to that of control (Fig S5 K, N, P). Also, Capu-GFP did not localise at the basket (Fig S5Q-Q’). Therefore, the role of Fhos and Capu in basket assembly was ruled out. Overall, on the basis of the genetic perturbations, the roles of formins in spermatid maturation can be categorised as-affecting NBs but not ICs (Dia and DAAM), affecting both NBs and ICs (Frl and Form 3), and affecting neither NBs nor ICs (Fhos and Capu).

Altogether, the genetic analysis suggested that the basket is exclusively formed through Dia and DAAM. Consistent with this conjecture, we found a visible enrichment of Dia at the basket region (arrowhead, Fig 4K-K’). Therefore, we concluded that the basket consists of contractile actin-membrane scaffold formed by Rho1, Dia and DAAM activities.

### N-BAR protein, Amphiphysin, recruits Rho1 and activates the F-actin assembly at the basket domain

Local tension and curvature of the membrane bilayer triggers F-actin assembly (Aspenstrom, 2014; Yang and Saif, 2007). TEM sections revealed that extensively folded membrane surrounds the spermatid heads (Fig 1E-G), and time-lapse observations after the laser ablation (Fig 2B) suggested that the spermatid heads continuously press onto the HCC. Hence, mechanical tension along the HCC plasma membrane could initiate the F-actin assembly forming the basket domain. Association of the Clathrin-family of coat protein complex bends and stabilises the membrane curvature, and promotes endocytosis (Hinrichsen et al., 2006). Also, the recruitment of BAR domain proteins onto negatively-charged membrane buds stabilises the membrane curvature and activates the BAR domain proteins, which sequester specific Rho-GTPases (Aspenstrom, 2014).

Expression of the GFP-tagged Clathrin-light-chain (*PpY>eGFP-Clc*, Fig 5A) and the BAR domain protein Syndapin (*PpY>Synd-GFP*, Fig 5B-B’) in the HCC marked the actin cap. The eGFP-Clc expression surrounded the basket and the caplet domain (arrowhead and arrow, Fig 5A-A’), whereas Synd-GFP co-localised with F-actin of the basket domain (arrows, Fig 5B-B’). Synd associates with the curved membrane through the SH3 and the BAR-domain. The negatively charged Phosphatidylinositol 4, 5-bisphosphate (PIP_2_) is essential for Syndapin binding (Takeda et al., 2013). Overexpression of Synd-SH3-eYFP (Fig S6A-A’) marked the basket, whereas that of the Synd-R129E R130E-eYFP, carrying mutations in the PIP_2_ interacting residues of the BAR domain, failed to mark the basket (Fig S6B-B’). This result further indicated that the basket domain is rich in the negatively charged membrane, particularly PIP_2_. Therefore, to understand the role of membrane remodelling proteins in this process, we performed a limited RNAi screen.

**Figure 5.**
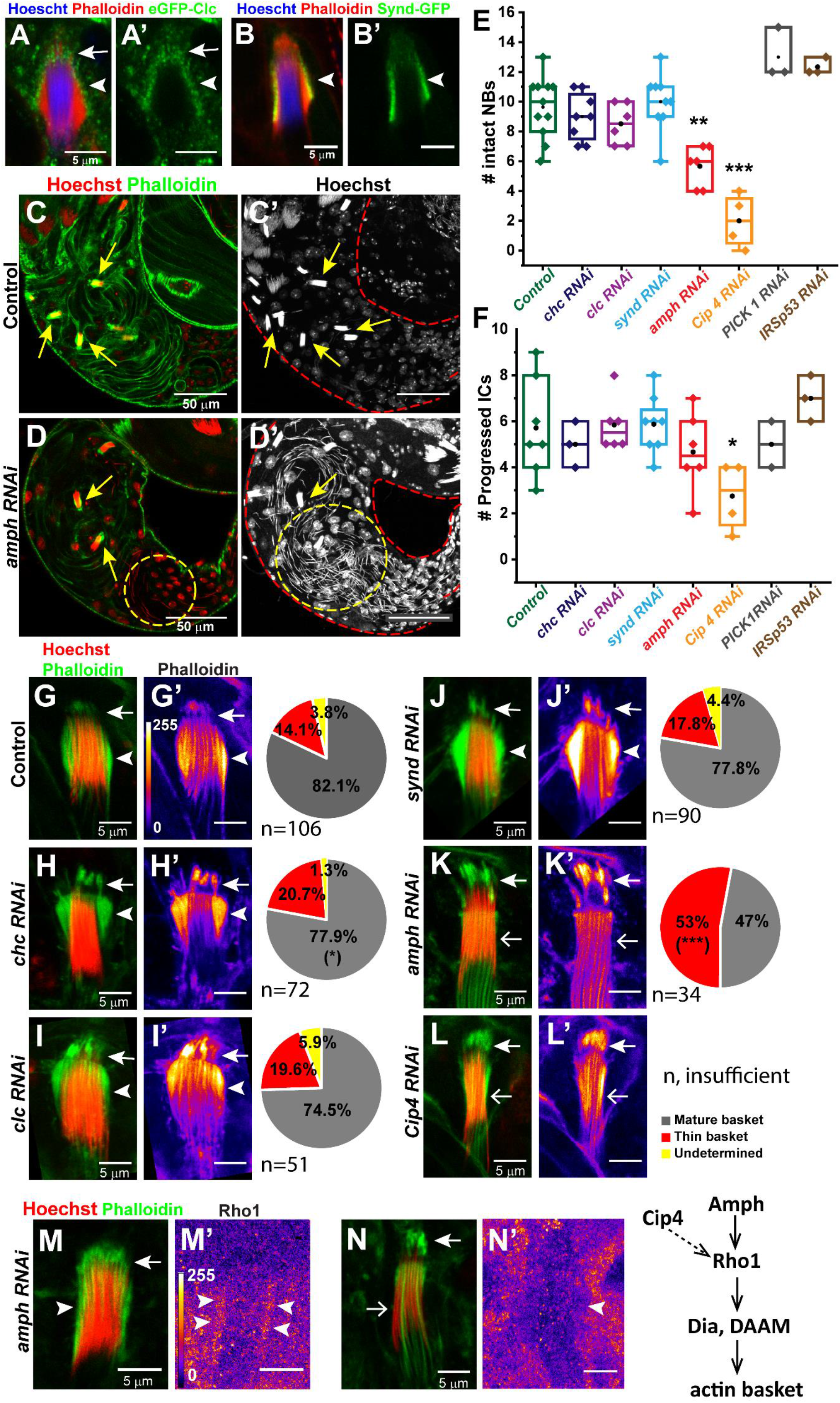
The BAR domain proteins, Amphiphysin and Cip4, are required to assemble the basket domain. **A-B)** Clathrin-Light-Chain (*PpY>e-GFP clc)* (A-A’), and Syndapin (*PpY> synd-GFP*) (B-B’) localisations around and in the basket domain (arrowheads). **C-D)** Confocal images of TE of Control (C-C’) and *amph RNAi* (D-D’) testes, stained with Hoechst (red/ white) and Phalloidin (green). C’, D’ represent maximum intensity projections of the Hoechst channel, depicting all the NBs in the testes. Yellow arrows mark intact testes, and yellow dashed circle marks disrupted NBs. **E-F)** Box and dot plots depicting the number of intact NBs (E) and the progressed ICs (F) in the testes expressing different RNAi constructs in the cyst cells. The pairwise significance of difference with respect to the control value was estimated using Mann-Whitney U tests, and the p-values-* <0.05, ** <0.01, *** < 0.001 are indicated on each box. **G-L)** The morphology of the caplet (arrows) and the basket (arrowheads) domains in different RNAi backgrounds, stained with the Hoechst dye (red) and Phalloidin (green/ FIRE). F-actin levels are depicted in a false colour heat map (FIRE) in the right panels, and the associated pie charts depict the penetrance of the phenotypes in these backgrounds. Open arrows indicate actin sleeves around the spermatid tails. **M-N)** Rho1 localisation (FIRE, M’, N’) in the *amph RNAi* background, stained with the Hoechst dye (red) and Phalloidin (green). Arrows indicate the caplets, arrowheads indicate the basket, and the open arrow indicates the actin sleeve, respectively.

Clathrin consists of three heavy chains, each associated with a light chain. This triskelion arrangement forms a cage-like lattice on the membrane (Kirchhausen et al., 2014). The clathrin heavy chain (chc) and light chain (clc) RNAis did not alter the number of intact NBs in the TE (Fig 5E; S6E, G), as well as the progressed ICs (Fig 5F; S6F, H). However, we noticed a small but significant decrease in the number of mature baskets in the *chc RNAi* background (p=0.040, pie chart, Fig 5H). A similar reduction was seen in *clc RNAi* testes, but the difference was not significant (pie chart, Fig 5I). Hence, we concluded that though Clathrin may control the membrane flux in the basket region, it is unlikely to have a central role in the basket assembly.

Amongst the BAR domain proteins, Synd (de Kreuk et al., 2011), Amphiphysin (Amph) (Yamada et al., 2009), Cdc42 interacting protein (Cip4) (Fricke et al., 2009), Insulin Receptor Substrate 53 kDa (IRSp53) (Goh et al., 2012) and Protein Interacting with C Kinase 1 (PICK1) (Rocca et al., 2008) are implicated in actin polymerisation. Synd/PASCIN overexpression recruited Dia at the pseudocleavage furrows of *Drosophila* embryos (Sherlekar and Rikhy, 2016); Cip4 binding antagonised Dia-induced actin polymerisation (Yan et al., 2013); and IRSp53 binds to mDia during filopodia extension (Goh et al., 2012). PICK1, on the other hand, is indicated to inhibit Arp2/3 (Rocca et al., 2008), and recruit F-actin to the site of endocytosis through myosin (Madasu et al., 2015) in different contexts. Since Dia, and potentially DAAM, are required for basket actin polymerisation, we tested the role of all these BAR domain proteins in the basket assembly.

The *Synd, IRSp53* and *PICK1 RNAis* did not affect the number of intact NBs (Fig 5E; Fig S6I, N, Q) and the IC progression (Fig 5F; Fig S6J, O, R). The basket morphology of *synd RNAi* testes was also not significantly different from that of the control (Fig 5J-J’; pie chart). Similar results were obtained due to the overexpression of the dominant-negative Synd-R129E R130E-eYFP (Fig S6C-D). Together, these data ruled out a role of Synd/PASCIN, IRSp53 and PICK1 in the basket assembly.

In *amph RNAi* testes, there was a significant reduction in intact NBs (Fig 5E, K-K’) with a large proportion of them having relatively thin baskets or only actin sleeves (open arrow and pie chart, Fig 5K-K’). The IC progression was not affected (Fig 5F; Fig S6K). The defect was even more severe in the *Cip4 RNAi* testes (Fig 5E, L-L’; Fig S6L). However, *Cip4 RNAi* also significantly reduced the number of progressed ICs (Fig 5F; Fig S6M), suggesting that the loss of NB could be contributed by the disruption of the upstream differentiation process.

Finally, in the *amph RNAi* background the Rho1 enrichment was eliminated from the thin baskets (Fig 5N-N’). This observation is consistent with a recent study which suggested that Rvs 167, the budding yeast homologue of Amph, is essential for the localisation of Rho1, in both a Rho GEF dependent and independent manner during cytokinesis (Cundell and Price, 2014). Rho1 accumulates at the cytokinetic furrow in a PIP_2_-dependent manner (Yoshida et al., 2009). Furthermore, the association of the BAR domain proteins stabilises phosphoinositide domains (Zhao et al., 2013). Hence, we concluded that Amph and possibly the PIP_2_ recruitment and stabilisation on the HCC membrane surrounding the spermatid heads could recruit and activate Rho1, and promote the F-actin assembly, as well as the basket morphogenesis.

### Non-muscle myosin-II maintains the F-actin structure and the integrity of the basket domain

Time-lapse imaging revealed that the basket is a contractile structure (Fig 2A). In addition to Formins, the GTP-bound Rho activates a serine/threonine kinase called Rho kinase (Rok/ROCK), which regulates several cytoskeletal rearrangements by phosphorylating a large number of target proteins (Amano et al., 2010). The Rok-GFP expression under the regulatory myosin-light-chain (*spaghetti-squash*) promoter (*sqh::Rok-GFP*) highlighted Rok enrichment in the basket domain (arrowhead, Fig 6A-A’). One of the downstream targets of Rok is Spaghetti-squash (Sqh), the *Drosophila* orthologue of the regulatory light chain (RLC) of the non-muscle-myosin-II (NM2) (www.flybase.org). NM2 exists in an auto-inhibited folded confirmation. The phosphorylation at Thr 20 and/or Ser 21 of Sqh/RLC relieves the heavy-chain (HC) (Zipper/ Zip in *Drosophila*) auto-inhibition and leads to oligomerisation of NM2 into bipolar filaments. Active NM2 oligomers crosslink F-actin into antiparallel bundles and generate contractile force along the filament (Liu et al., 2017). NM2 drives morphogenetic movements, such as dorsal closure, cytokinesis, and wound healing (Franke et al., 2010). We found that both Zip/HC and Sqh/RLC localise at the basket in the *PpY>zip-GFP* (Fig 6B-B’) and *sqh::sqh-GFP* (Fig 6C-C’) backgrounds. Also, phosphorylated Sqh/RLC (p-MLC) immunostaining exclusively labelled the basket domain (Fig 6D-D’). Thus, Rho1-dependent recruitment and activation of Rok are likely to promote the NM2 association with the basket.

**Figure 6.**
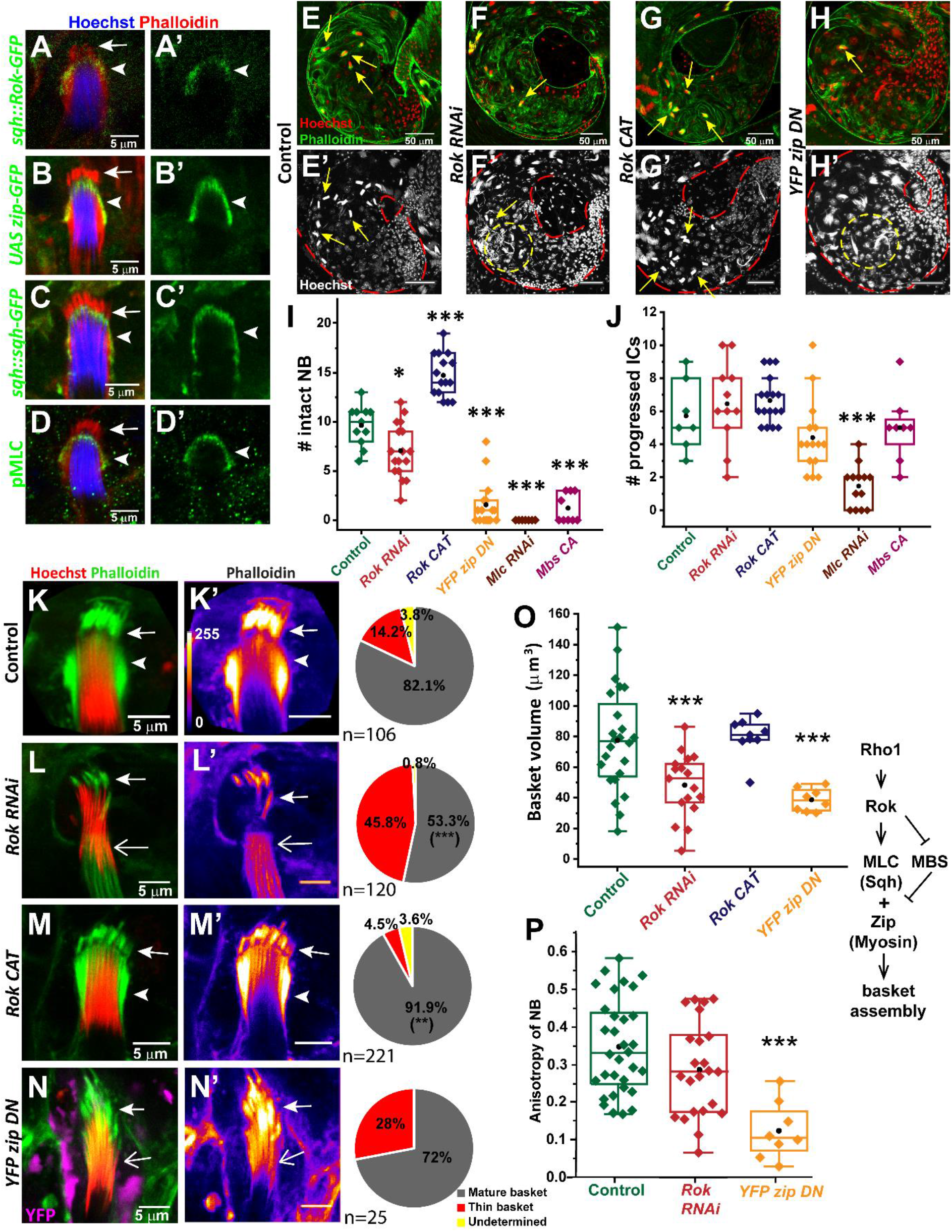
Rok and nonmuscle myosin-II (NM2/Zipper) function is essential for the basket assembly. **A-D)** NB from testes expressing (green) *Sqh*::*Rok-GFP* (A-A’), *PpY>zip-GFP* (B-B’), *sqh::sqh-GFP* (C-C’), and stained with anti-pMLC (D-D’), respectively, along with Hoechst dye (blue) and Phalloidin (red). Note that the localisation pattern is restricted to the basket (arrowheads) domain, and is absent from the caplets (arrows). **E-H)** TE regions of Control (E-E’), *PpY>Rok RNAi* (F-F’), *PpY> Rok-CAT* (G-G’), and *PpY>YFP-zip DN* (H-H’) testes, stained with Hoechst (red/ white), and Phalloidin (green). **I-J)** Box plots depict the number of intact NBs (I) and the number of progressed ICs (J) in different genetic backgrounds. The pairwise significance of difference with respect to the control value was estimated using Mann-Whitney U tests, and the p-values-* <0.05, ** <0.01, *** < 0.001 are indicated on each box. **K-N)** High magnification images depict the morphology of the actin cap in Control (K-K’), *Rok RNAi* (L-L’), *Rok CAT* (M-M’) and *YFP-zip DN* (N-N’) backgrounds, stained with Hoechst (red) and Phalloidin (green/FIRE). Arrows indicate the caplet, arrowheads indicate basket, and the open arrow indicates the actin sleeves around the spermatids. The F-actin levels are shown in false colour (K’-N’) intensity heat map according to the LUT on K’. Pie charts depict the frequency of different basket phenotypes. The significance was calculated using the Fisher’s exact test, and the p-values-* <0.05, ** <0.01, *** < 0.001 are indicated on each box. **O-P)** Box plots depict the volume of the basket domain (O) and the organisation (Anisotropy) of the spermatid head alignment in NBs (P) in the Control, *Rok RNAi* and *YFP-zip DN* testes, respectively. Significance was calculated using One-way Anova®, and the p-values-* <0.05, ** <0.01, *** < 0.001 are indicated on each box.

In addition to the contractile behaviour, the laser ablation study suggested that basket maintains the spermatid heads in a tight bundle (Fig 2B). Therefore, to understand whether an actomyosin contraction is responsible for bundling the spermatid heads together, we perturbed the functions of the Rok and NM2 complex in cyst cells. Expression of the *Rok RNAi* (Fig 6F-F’) and the dominant-negative *YFP-zip DN* transgene, lacking the actin-binding motor domain (Fig 6H-H’) by *PpY-Gal4* caused abnormal accumulation of disrupted spermatid bundles in the TE. These perturbations also reduced the average NBs significantly (Fig 6I) without affecting the progressed ICs (Fig 6J; S7A, C). Also, *sqh* knockdown (*Mlc RNAi*) severely affected both NB integrity and IC progression (Fig 6I-J; S7D-E). As expected, the overexpression of the constitutive-active Rok catalytic domain (*Rok-CAT*) selectively increased the NB accumulation (Fig 6I), without affecting ICs (Fig 6J; S7B). We also noted that YFP-zip DN failed to localise at the basket domain (magenta, Fig 6N). Altogether, these results indicated that NM2 localises to the basket directly through its motor domain, and the actomyosin activity is essential for either the basket assembly or maintenance or both.

In the *Rok RNAi* testes, more than 45% of the NBs were associated with thin or deformed baskets (pie chart, Fig 6L). The average volume of the basket domain was significantly reduced (Fig 6O). In contrast, more than 90% of the NBs in the *Rok-CAT* overexpression background had mature baskets (pie chart, Fig 6M), with very similar volume to the control (Fig 6O). The similarity of the results obtained from the Rho1CA (Fig 3) and Rok-CAT overexpression backgrounds suggest that upregulation of the Rho-Rok pathway may stabilise the basket through the NM2 activation, which would reduce the frequency of sperm release causing accumulation of NBs at the testicular base.

In both *Rok RNAi* and *YFP-zip DN* expressing testes, the proportion of mature baskets (Fig 6L, N), the basket volume (Fig 6O), and the anisotropy of NB alignment (Fig 6P) were reduced. These results suggested that the NM2 activity is essential for maintaining the F-actin level and the function of the basket required for gathering the spermatid heads.

Regulation of the myosin activity is vital to prevent over-constriction of the actomyosin cytoskeleton. The activity of myosin is maintained by reversible phosphorylation of Sqh/RLC through the regulatory Myosin-binding-subunit (Mbs) protein (Mizuno et al., 2002) and the phosphatase - Flapwing (Flw) (Kirchner et al., 2007). Mbs is inactivated through phosphorylation by Rok, which inhibits the dephosphorylation of Sqh/RLC by Flw. Expression of the constitutively-active Mbs-N300, lacking the N-terminal Rok phosphorylation site, reduced the number of intact NBs significantly without affecting the IC progression (Fig 6I-J; S7F, G) which is consistent with the above conclusion. However, contrary to our expectations, *Mbs* and *flw RNAis* had minimal effect on the NB accumulation (Fig S8A), basket volume (Fig S8C) and basket morphology (Fig S8D, E). These results indicate that Mbs and Flw are unlikely to regulate the NM2/RLC function at the basket. Also, the overexpression of catalytically inactive Rok (Rok CAT-KG), the constitutively-active Sqh-EE and the dominant-negative Sqh-AA, lacking both the phosphorylation sites, had no significant impact on NB count and basket volume (Fig S8B and C). We reasoned that the level of Sqh-AA and Sqh-EE expression were inadequate for interfering with the NM2 function at the basket.

Together, these results suggested that Rok-mediated activation of NM2 is essential for maintaining the basket domain, and the actomyosin contractility keeps the NB together. Further, we hypothesised that the NM2 activity would also be essential for maintaining the actin-membrane scaffold in the basket region.

### Rok dynamically maintain the NM2 localisation and F-actin organisation in the basket domain

To test these conjectures, we inhibited Rok activity by treating testis expressing either Sqh-GFP or Utr-GFP with Rockout™, an ATP competitive inhibitor of Rok (Yarrow et al., 2005). Treatment with 200 µM Rockout for one hour significantly reduced the number of intact NBs (Fig 7A-C) and accumulated disrupted spermatids within the TE (yellow dashed circle, Fig 7B). Although the DMSO treatment also caused some abnormality, (arrowheads, Fig 7A-A’), the defects were severely enhanced upon Rockout treatment (Fig 7B-B’). It also significantly disrupted the basket morphology (Fig 7D, E). Depth coding indicated that NBs and associated actin caps remain within a compact volume inside the HCC in control (Fig 7D’-D”). In contrast, most of the NBs appeared disrupted and spread over a large volume after the Rockout treatment (Fig 7E’, E’’). The treatment also eliminated Sqh-GFP enrichment from the basket (Fig 7D’’’, E’’’, and F)

**Figure 7.**
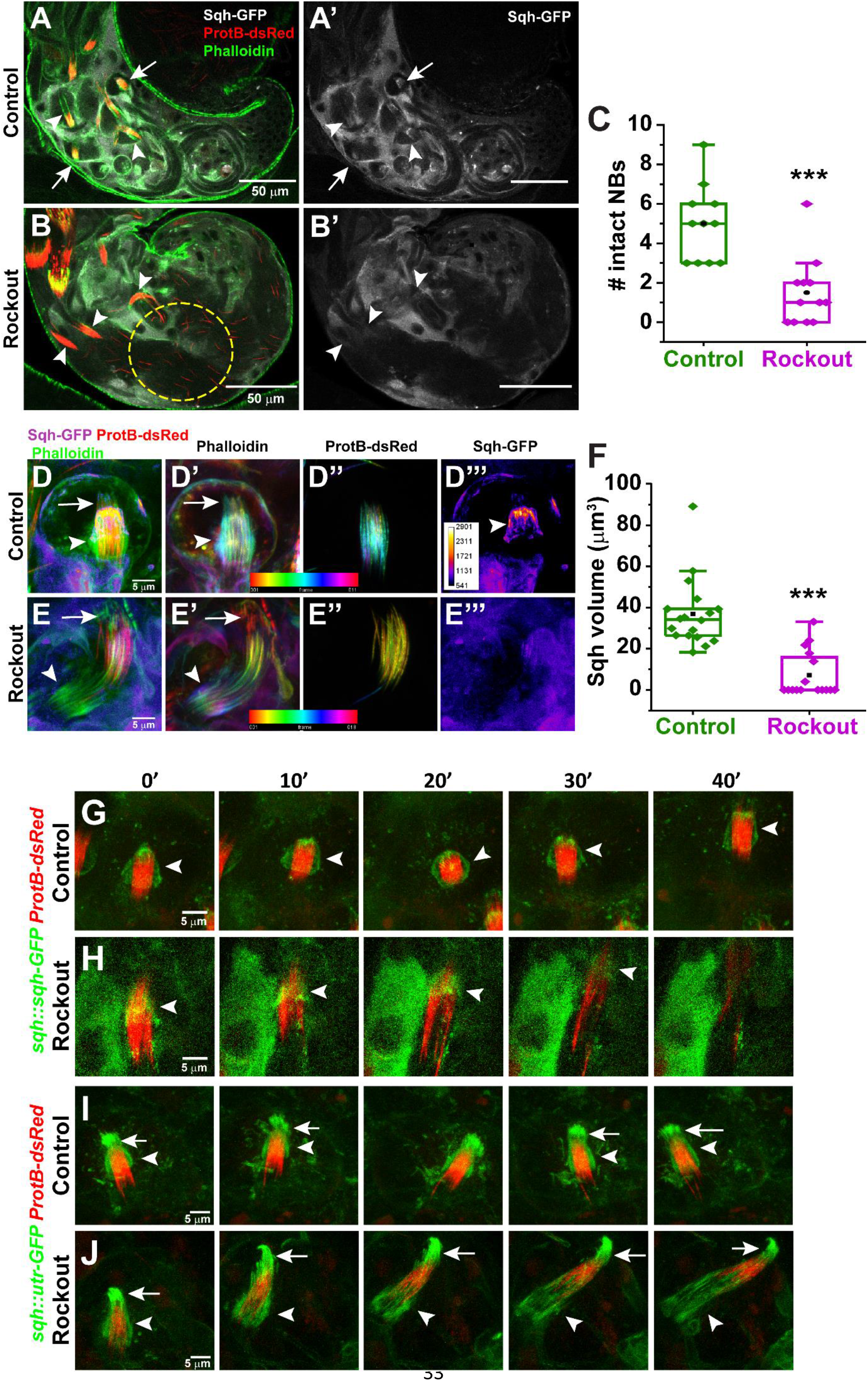
Rok-dependent NM2 recruitment at the basket is essential for compacting the spermatid heads into a tight bundle. **A-B)** Confocal images from *sqh::sqh-GFP* (white) and *ProtB-dsRed* (red) expressing testes, treated with 2% DMSO (Control, A-A’) and 200 µM Rockout (B-B’), for one hour and stained with Phalloidin (green). Arrows mark intact bundles, arrowheads mark bundles which have been perturbed by DMSO/ Rockout treatment. Yellow dashed circle marks the disrupted sperm heads. **C**) The number of intact NBs in control and Rockout treated testes. Pair-wise significance of difference was estimated using Mann-Whitney U test, and the p-values-* <0.05, ** <0.01, *** < 0.001 are indicated on the box. **D-E)** High magnification images of NBs from Control (D-D’’’) and Rockout treated testes (E-E’’’). D, E represents the merged images, depicting Sqh-GFP (FIRE), ProtB-dsRed (red) and Phalloidin (green). Arrows mark caplet, and arrowhead marks basket. D-D’’ and E-E’’ represent depth coded Z-stack for Phalloidin (D’, E’) and ProtB-dsRed (D’’, E’’) in Control (D’-D’’) and Rockout treated (E’-E’’) testes, respectively. The colour code is according to the scale depicted in the images. The Fire LUT (E”‘) represents the intensity heat map. **F)** Basket volume marked by Sqh-GFP in Control and Rockout treatment testes. Significance was calculated using Mann-Whitney U test, and the p-values-* <0.05, ** <0.01, *** < 0.001 are indicated on the box. **G-J)** Time-lapse images of testes expressing *sqh::sqh-GFP* and *ProtB-dsRed* (G, H), and *sqh::utr-GFP* and *ProtB-dsRed* (I, J), respectively, after the DMSO (Control) and Rockout treatments. The arrows indicate caplet and the arrowheads indicate the basket domains.

The time-lapse imaging further suggested that the Rockout treatment dispersed the Sqh-GFP enrichment around the NB in 30 minutes, and concomitantly the spermatid heads were dissociated from each other (Fig 7H; Video S8), which was not seen upon the DMSO treatment (Fig 7G and I; Video S7 and S9). The treatment also disrupted the basket marked by Utr-GFP in 20 minutes (arrowhead, Fig 7J; Video S10). Together these observations suggested that the Rok-dependent NM2 activation localises the myosin motor onto the basket, which maintains the basket integrity and applies a compacting force on the spermatid heads, to keep them in a parallel bundle. However, the Utr-GFP did not disperse from the basket region upon the Rockout treatment. Hence, we concluded that Rok and NM2 are not essential for the F-actin assembly at the basket.

Altogether, the above results suggest that mechanical tension on the somatic cyst cell membrane and membrane curvature induced by the intruding spermatid heads could recruit and activate specific BAR domain proteins, Amphiphysin and perhaps Cip4. The recruitment of BAR domain proteins could further recruit and activate Rho1, triggering the F-actin assembly through the formins, Dia and DAAM. Rho1 also recruits Rok, which activates NM2. Active NM2 crosslinks the actin filaments at the membrane and exerts a lateral clamp-like contractile force gathering the spermatid heads into a tight parallel bundle (Fig 8). The membrane-actomyosin scaffold also resists further intrusion by the spermatid heads.

**Figure 8.**
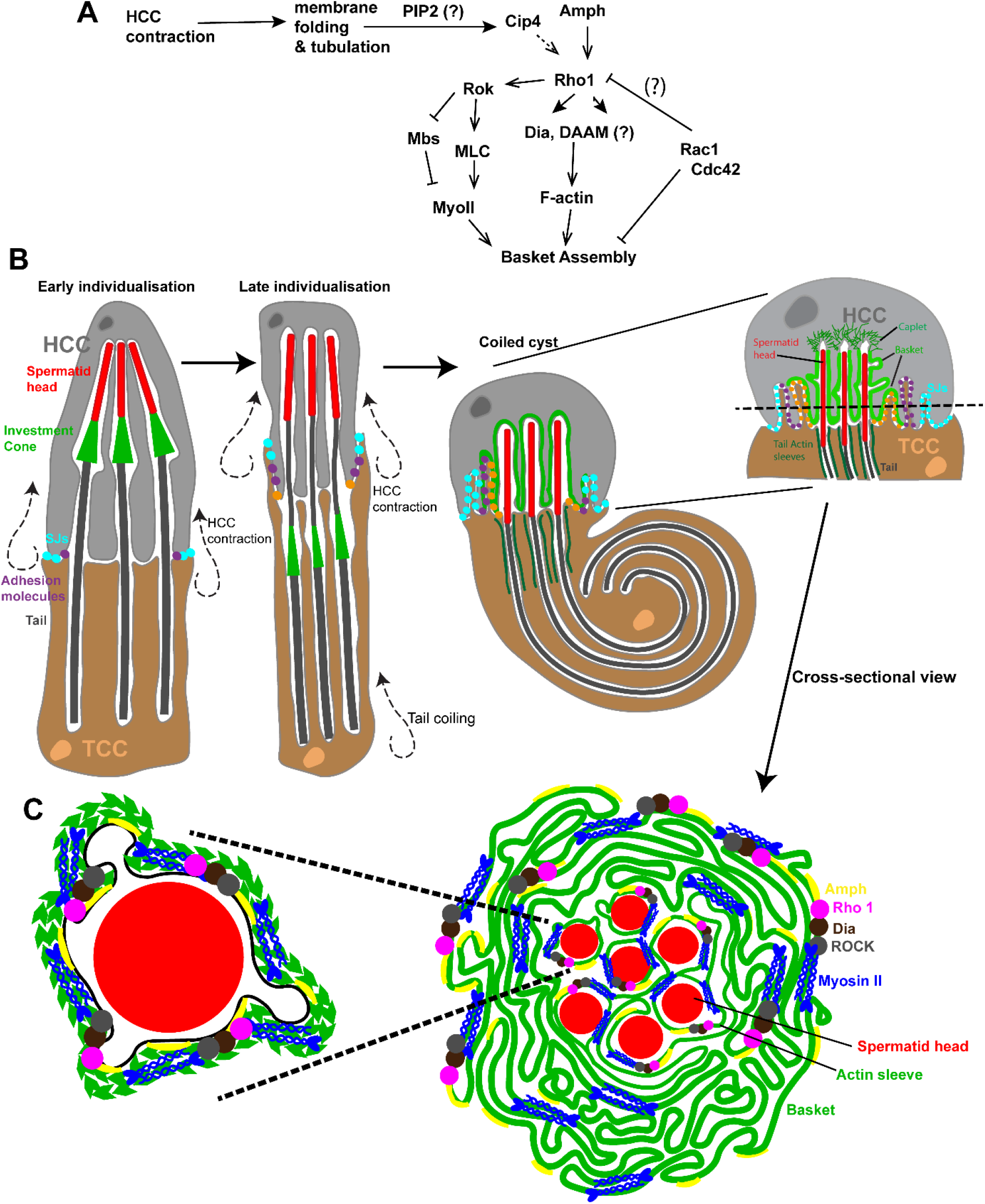
Assembly of the contractile actomyosin scaffold around mature spermatid heads butting into the somatic head cyst cell. Flow chart and schematic depict how remodelling of the membrane during individualisation to coiling transition of the cyst, can potentially recruit Amph, which may intern recruit Rho 1. Rho 1 forms the basket and maintains NB arrangement, via the activation of the formins, Dia and DAAM, and the Rho kinase Rok, which in turn activates NM-2. (Refer to Fig. 1E for cross-section morphology depicted in C)

## Discussion

Membrane indentation is the first step of inducing mechanical damage to the plasma membrane. Artificial deformation or indentation of tissue-cultured cells induces actin accumulation at the site of indentation (Lou et al., 2019; Yang and Saif, 2007). Hence, above results provides the first insight into the events initiated at the start of membrane puncturing completely *in vivo*. Also, preliminary assessment suggested that the membrane-associated with the basket domain is rich in PI(4,5)P_2_, which is essential for the interaction with the BAR domain. Therefore, the spermatid head intrusion is also likely to trigger local alteration of lipid metabolism at the HCC plasma membrane. The mechanism that initiates the recruitment of specific BAR domain proteins to the membrane is unclear. These results suggest that the basket domain would be a useful paradigm to investigate how the membrane tension could alter membrane composition and promote specific recruitment of the BAR domain proteins, particularly Amphiphysin and potentially Cip4, at the HCC plasma membrane.

The BAR domain proteins are shown to interact with the Cdc42 and Rac1 family of Rho-GTPases and nucleate F-actin assembly through the WASP/WAVE NPFs (Fricke et al., 2009). A Cip4-like protein, Cdc15/FBP17, is also shown to associate with Dia and DAAM, during the filopodia assembly (Aspenstrom et al., 2006). The consequence of the interaction is still unclear. A separate study further suggested that Cip4 antagonises Dia during the extension of membrane furrows in *Drosophila* embryos (Yan et al., 2013). The genetic analysis suggests that the BAR domain proteins, Amphiphysin and potentially Cip4, could initiate the F-actin assembly at the basket domain in the HCC, exclusively through a Rho-Dia/DAAM-mediated process. The loss of Amph eliminated Rho1 recruitment at the basket domain, as well as F-actin assembly. Thus, the results provide an indication of Amphiphysin-mediated F-actin assembly through Dia and DAAM *in vivo*. Further studies would be required to confirm and understand the molecular basis of this interaction.

Above results, together with the earlier report (Dubey et al., 2016), also depict that the HCC membrane curved around each invading spermatid head nucleates two distinct actin networks. The dendritic F-actin associated with caplets form due to WASp-Arp2/3 recruitment and activation in response to the inadvertent invasions, or spermatid head-butting, and is a somewhat transient event. It exerts a counterforce along the direction of invasion to repel every intrusion (Dubey et al., 2016). The F-actin at the basket, nucleated by the Rho1-Dia/DAAM pathway, downstream to Amphiphysin, is relatively stable and applies a lateral contractile force due to actomyosin activity to bundle the spermatid heads together, like a clamp. The genetic and pharmacological analyses show that these two actin networks form independently on two mutually exclusive and radially segregated membrane domains. They also form at different time scales. The caplets are transient and form within minutes of the invasion. Hence, they indicate a cellular response to rapid change in membrane tension. The baskets, on the other hand, grows slowly after individualisation and matures after coiling.

The context of the basket assembly provides a clue to the underlying trigger. The HCC and TCC are equally extended during the individualisation stage with their respective plasma membranes tightly stretched around the individual spermatids (Tokuyasu et al., 1972a). Post individualisation, the cysts proceed for coiling, and the HCC contracts (Dubey et al., 2019; Tokuyasu et al., 1972b), which could loosen and fold the excess membrane around spermatid heads triggering the recruitment of Amphiphysin (Fig 8). Therefore, the basket connotes to a response generated due to a change of membrane tension over time. We also find that the functions of Rho1 and Rac1 are mutually antagonistic at the basket domain, and Rac1 may also have a negative influence on the caplet and tubule dynamics. Thus, the paradigm provides an opportunity to tease out how cells sense, discriminate and respond to stress dynamics in the membrane.

Together with the previous report (Dubey et al., 2016), it is clear that the basket and caplets are composed by distinct F-actin networks. A similar juxtaposition of different F-actin was reported at the immunological synapse in T-cells, and during wound healing in *Drosophila*. At the immunological synapse, three types of actin networks - branched, linear and focal clusters - are arranged in a concentric manner (Hammer et al., 2019). Each of these structures presumably helps in the clustering of proteins such as T-cell receptors. As a result, disruption of the network leads to defects in receptor clustering (Murugesan et al., 2016). During the wound closure in *Drosophila* embryo, F-actin networks formed by the Rho1, Rac1, Cdc42 GTPases surround the wound site (Abreu-Blanco et al., 2014). Although Annexin and prepatterning of RhoGEFs were implicated in the Rho GTPases recruitment (Nakamura et al., 2017), how three distinct Rho-GTPase domains assemble on the membrane is still uncertain. Also, since each of these actin networks is partially overlapping, and concentrically arranged, any perturbation to any one of them, affects the others (Abreu-Blanco et al., 2014). Hence, it is challenging to dissect specific roles and mechanisms of regulations.

In comparison, the actin domains on the HCC membrane encompassing the actin cap region are mutually exclusive and well separated. Two distinct mechanical actions trigger them, and we now demonstrated that one could independently perturb the basket and caplets. Therefore, it would be interesting to study how two different F-actin nucleation mechanisms are established on the HCC plasma membrane in real-time during sperm coiling and release in the future.

Finally, a majority of actomyosin structures are known to undergo constriction to achieve complete closure, such as during cytokinesis, wound healing and dorsal closure. Specific actomyosin arrangements also provide scaffolding or stabilising force. The actomyosin association with the adherens junctions at the apical boundary of epithelia serves to stabilise the contact site and helps to recruit the junctional components (Lecuit et al., 2011; Schwayer et al., 2016; Zenker et al., 2018). In all these instances the actomyosin complex is organised in a ring-like form on a 2D-plane. Here, we showed that a 3-D membrane-actomyosin assembly could generate lateral constricting force orthogonal to the direction of intrusive force, to clasp the spermatid heads together. This finding demonstrates a novel cell adhesion strategy, which is unprecedented. Further investigation is required to identify the physical basis of such an assembly and its functioning.

## Materials and Methods

### *Drosophila* stocks and culture

All *Drosophila* stocks and crosses were maintained in standard cornmeal media at 25 °C. Male progeny of *UAS-RNAi* crosses was collected and kept at 28 °C for 4 days, to achieve optimum Gal4 expression. All the stocks have been listed in Table S1. All experiments involving transgenic *Drosophila* lines were approved by the Institute Bio-Safety Committee, DBT, Govt. of India.

### Whole-mount immunostaining

Testes were dissected in 1x PBS and fixed for 30 mins in 4% paraformaldehyde (PFA), followed by 3 washes of 10 mins each, with 0.3% PTX (0.3% Triton X 100 in 1 x PBS). Immunostaining samples were then incubated in blocking buffer (2 mg/ ml Bovine Serum Albumin (BSA) in 0.3% PTX) for 30 mins, followed by overnight incubation in the primary antibody. Samples were then washed and incubated with Alexa-conjugated secondary antibodies (Invitrogen) for 2 hours, followed by incubation in 0.001% Hoechst 33342 (Sigma-Aldrich), and 10 μM FITC Phalloidin/ Atto 647 Phalloidin (Sigma-Aldrich) for 30 min. Samples were then washed and mounted with a drop of Vectorshield™ mounting medium on a glass slide. The primary antibodies used were anti-Fas III (1:100, DSHB), anti-Rho 1 (1:100, DSHB), anti-Dia (1:2500, courtesy Prof. S Wasserman, UCSD) and anti-pMLC (1:25, Cell Signalling Technologies).

### STED (Stimulated Emission Depletion)

For STED imaging, *ProtB-dsRed* testes were processed as described above and stained with Atto647-Phalloidin. The samples were mounted on glycerol, and #1.5 coverslips were used. Depletion for both fluorophores was done using a 775 nm laser. Time gating of 0.6<tg>6.6 was applied. Images were acquired in 2D STED mode.

### *Ex vivo* time-lapse imaging of testis

Live imaging was performed as described previously (Dubey et al., 2016). Images were acquired using either an Olympus FV3000 confocal laser scanning microscope with 60x, 1.42 NA oil objective, or Olympus FV1000 confocal laser scanning microscope with 60x, 1.35 NA oil objective. Rockout (Calbiochem) was used at a concentration of 200 µM. For FRAP, 1.4 µm^2^ area in the basket was bleached to 60% level using high intensity 488 nm laser for 2 seconds with 2 µs/pixel scan speed, and the recovery profile was recorded at 5.5-second/frame rate. The data were analysed using ImageJ® and Origin™.

### Laser ablation

Laser ablation was performed using the Spectra-Physics Mai Tai Ti-Sapphire laser system on Zeiss 710 microscope. For ablation, 408.2 mW of 800 nm laser was focussed in 0.75 µm^2^ (10×10 pixels) area for five iterations with a pixel dwell time of 12.6 µs/pixel.

### Image Analysis and Quantification

All images were analysed on ImageJ/Fiji® (ImageJ.net/Fiji). We used the Cell Counter™ Plugin for the NB and IC estimation and the FibrilTool Plugin (Boudaoud et al., 2014), which measures the orientation and anisotropy of fibrillary structures, to assess the parallel organisation of spermatid heads in each NB. A value of 1 indicates purely anisotropic (perfectly ordered or parallel), and 0 indicates disordered assembly. For FRAP, the intensities in the SUM projection was calculated from the FRAP ROI, as well as an arbitrary background ROI. The data was normalised and fitted to a single exponential using the Levenberg-Marquardt iteration algorithm in Origin™ 2019, based on which maximum recovery was calculated. The basket volume was estimated using Imaris™ (Bitplane AG). A volume filter of 100 voxels was applied, to eliminate stray F-actin puncta in the given ROI. For basket morphology analysis, there was a very small proportion of NBs for which we could not determine the basket morphology, due to technical reasons, such as if the NB was oriented perpendicular to the plane of imaging. These were categorized as ‘undetermined’.

Statistical analyses (One-way ANOVA, Mann-Whitney Test) were carried out using Origin™ 2019. Fisher’s exact test was performed using the Graphpad online calculator. For box and dot plots, ‘n’ values are indicated by the number of dots. For Fisher’s exact test, ‘n’ values are indicated near the pie chart or on the stacked bar graph.

### Transmission Electron Microscopy

TEM samples were prepared as per the previous description (Dubey et al., 2019).

## Glossary

TE: Terminal epithelium
HCC: Head cyst cell
TCC: Tail cyst cell
NB: Nuclei bundle/Spermatid head bundle
Caplet and Basket: parts of Actin cap
SJs: Septate Junctions
CC: coiled cyst

## Acknowledgements

We thank Richa Ricky, Benny Shilo, Steven Wasserman, Andrea Brand, John Belote, Vimlesh Kumar, Thomas Lecuit, and Eyal Schejter for generous sharing of reagents. We acknowledge the BDSC, VDRC, DSHB, IISER Pune fly facility and NCBS Fly Facility, India for providing fly stocks and antibodies. We also thank Lalit Borde for help with TEM, and Hina Mohammed for assistance with the Synd mutant experiment. We are grateful to Prof. Shilo for his continuous guidance and support.

## Authors’ contributions

TK and PD performed the experiments. TK and KR wrote the manuscript. SS provided the TEM data.

