## Supplemental Text and Figures for "Nanoscale indentation of plasma membrane establishes a contractile actomyosin scaffold through selective activation of the Amphiphysin-Rho1-Dia/DAAM and Rok pathway"

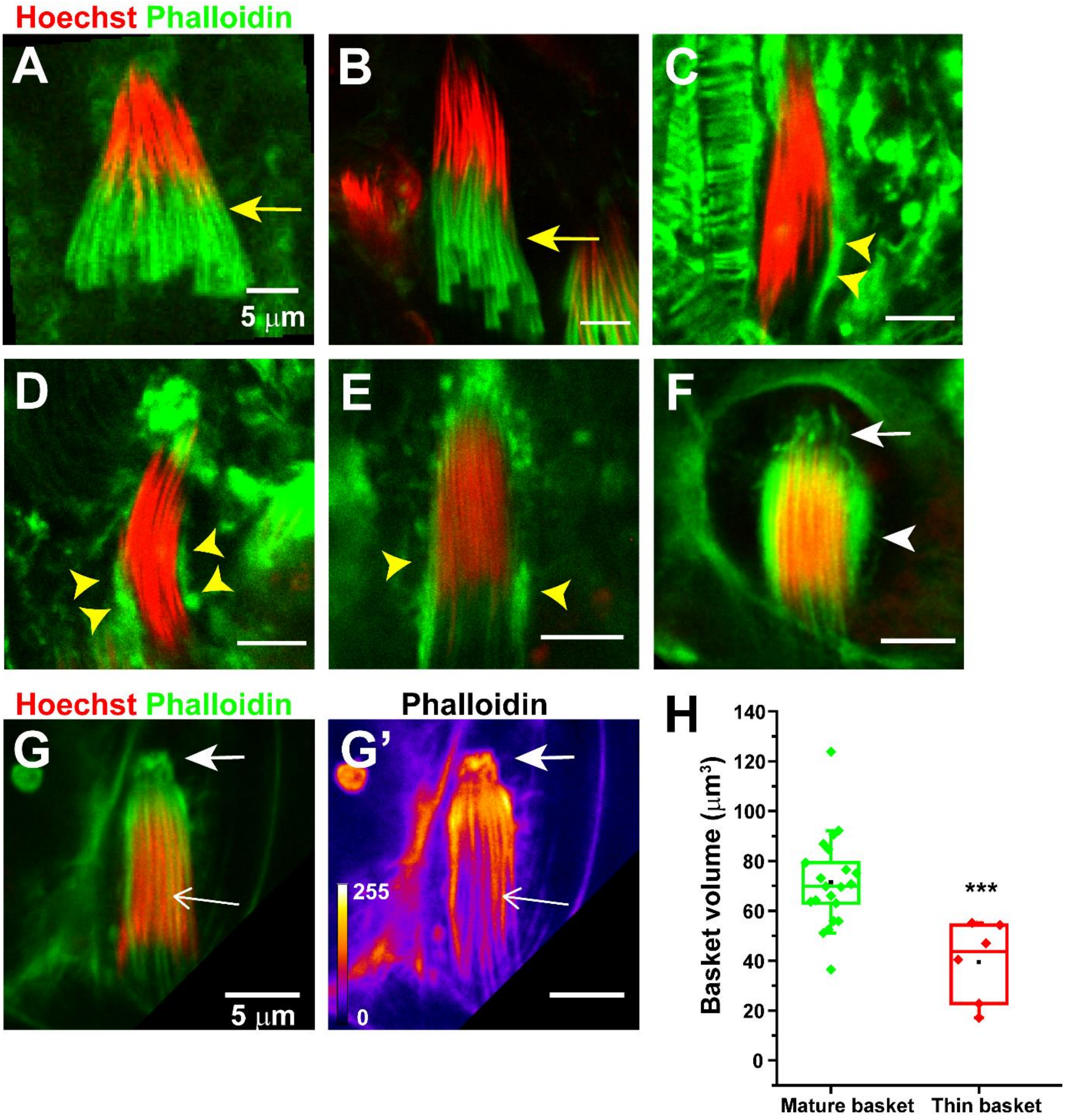


**Figure S1- Morphology of NB and associated actin structures through different stages of development**

**A-F)** The morphology of NB marked by Hoechst (red) and associated F-actin structure marked by Phalloidin (green) at different stages of maturation. At the early individualisation stage (A), the ICs (arrow, A) form around the caudal end of each spermatid head in a conical arrangement. As the ICs progress, the spermatid head bundle reorganises in a barrel-like form (B-E) and F-actin accumulates (yellow arrowheads, C-D) around the spermatid heads or nuclei bundle (NB). Post individualisation the spermatid bundle coils rapidly. During these stages (E-F), exclusively found within the TE region, the NBs are tightly compacted in a barrel form and are surrounded by the F-actin cap consisting of two morphologically distinct domains - the basket (arrowhead, F) and caplet domains (arrow, F).

**G)** Example of a 'thin' basket in WT testis, wherein mostly the actin sleeves are visible (open arrow). Arrows mark caplet. G' depicts phalloidin intensity according to the heat map depicted.

**H)** F-actin volume of mature and thin baskets. The pairwise significance of difference was estimated using One-way Anova®, p-value- *** < 0.001

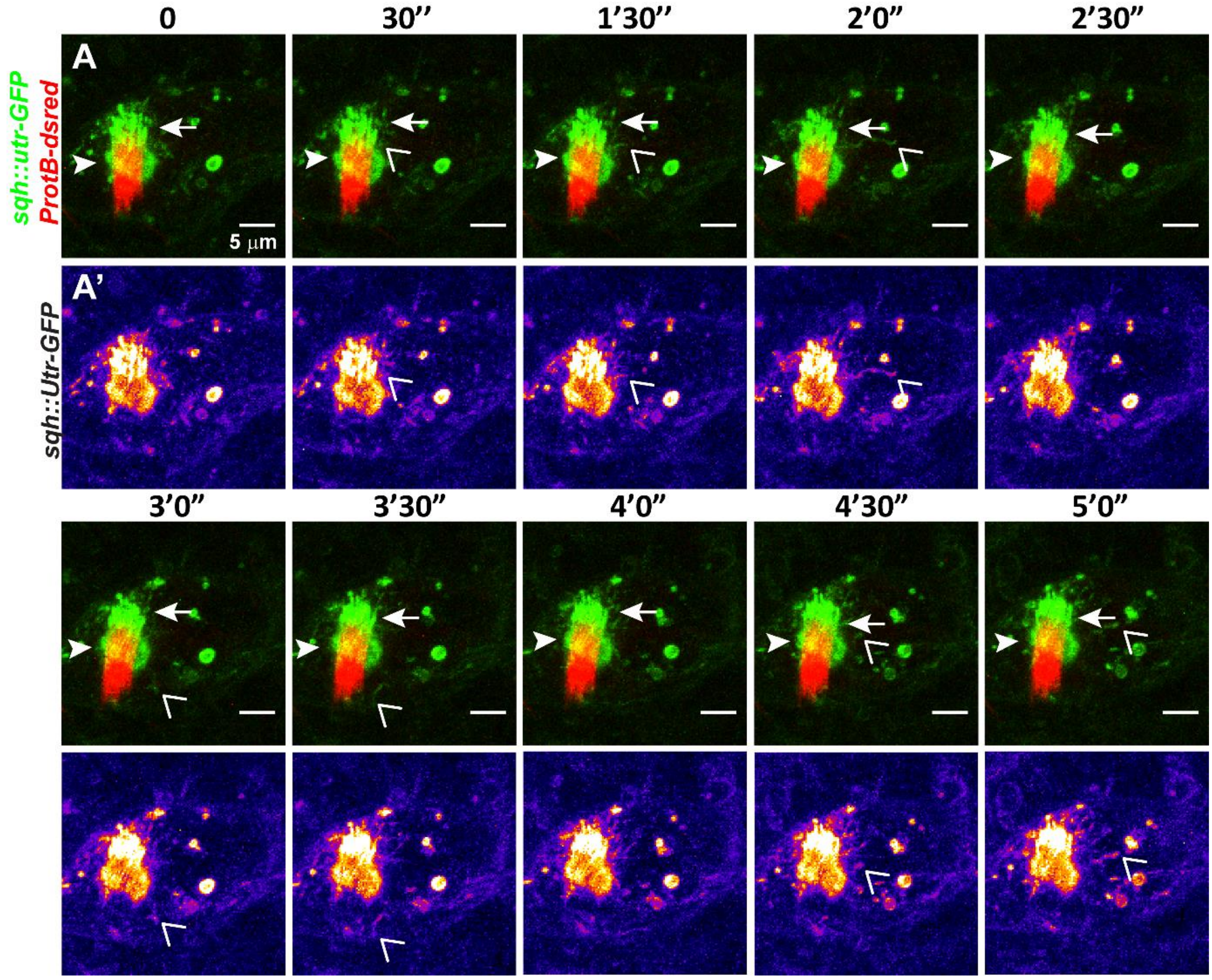


**Figure S2- F-actin-rich tubules periodically eject from the basket domain.**

Time-lapse images of a *sqh::utr-GFP* and *ProtB-dsRed* testis marking the NB (red) and actin cap (green/ FIRE). Arrows and arrowhead indicate the caplet and basket domains respectively, and open arrowheads indicate the tips of transient extension of actin tubules from the basket domain.

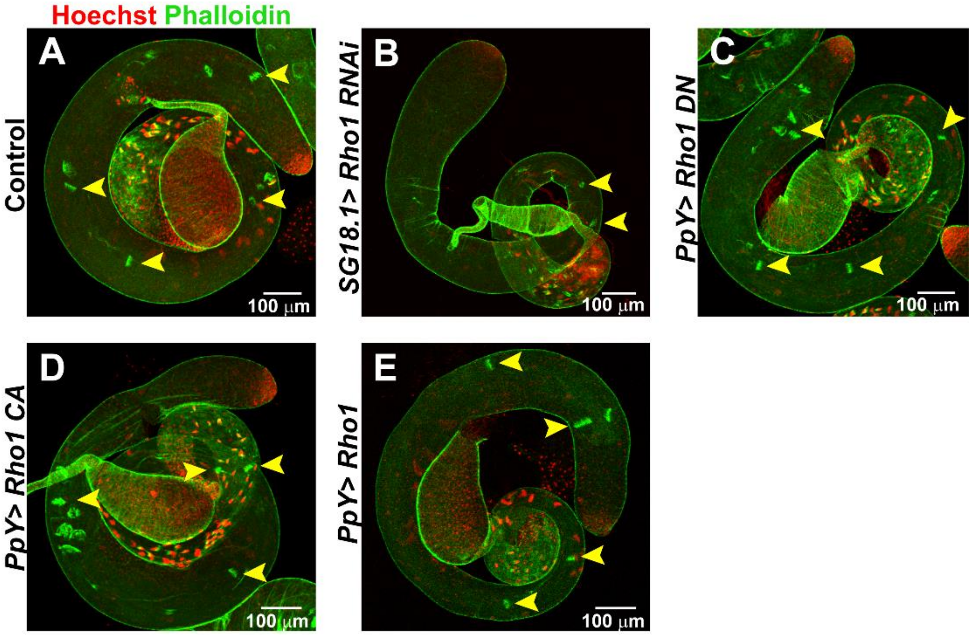


**Figure S3- IC progression is only affected in *SG18.1>Rho1 RNAi* testes**

**A-E)** Low magnification images of the entire testes of Control (A), *SG18.1> Rho1 RNAi* (B), *PpY> Rho1 DN* (C), *PpY> Rho1 CA* (D), and *PpY> Rho1* (E), stained with Hoechst (red) and Phalloidin (green). Yellow arrowheads mark progressed ICs.

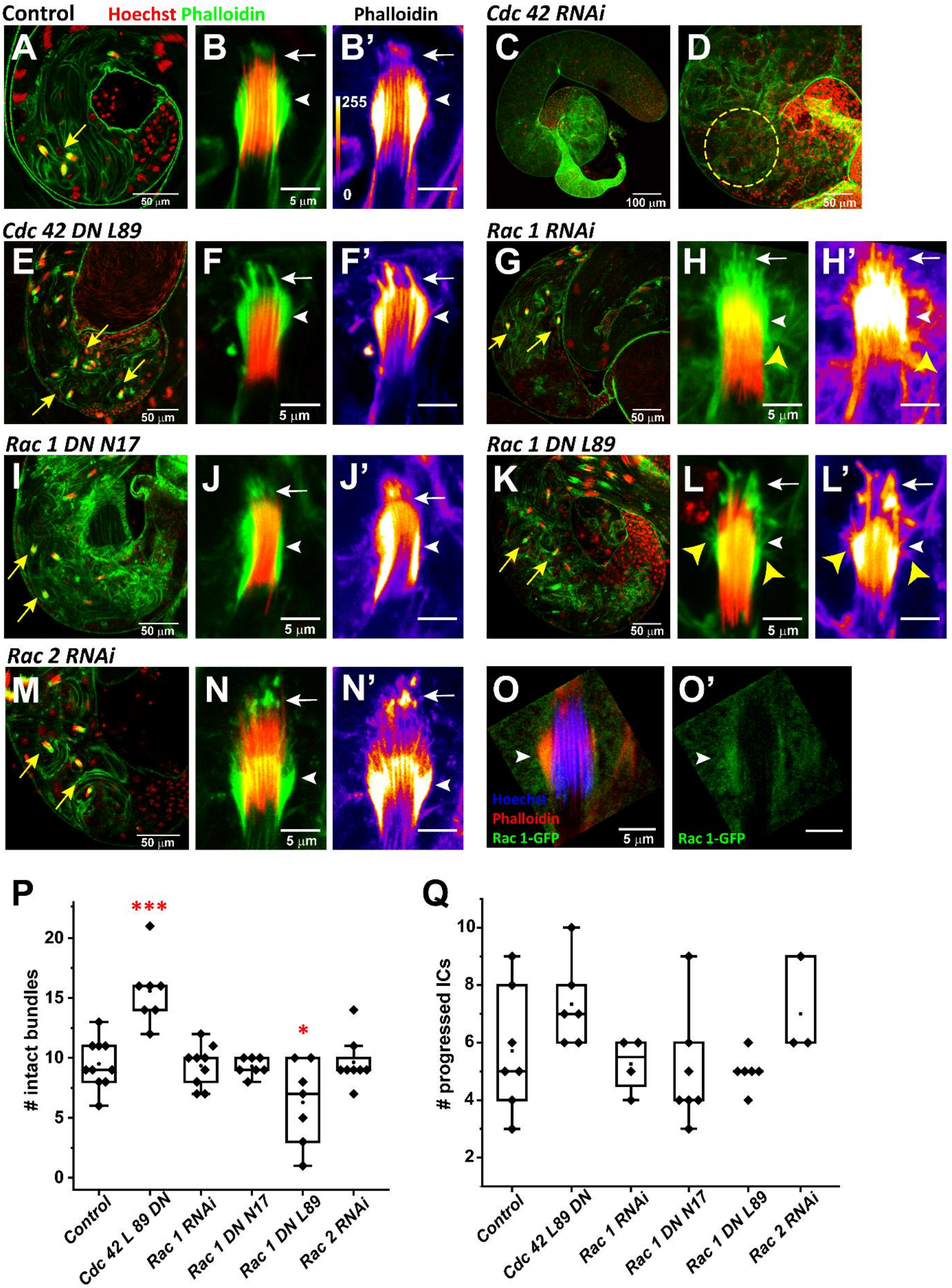


**Figure S4- Loss of Cdc42, and Rac1/2 in the cyst cells does not disrupt the basket morphology.**

**A-N)** Confocal sections of TE region (left column), and high magnification images of NBs associated actin caps (middle and right columns) from different genetic backgrounds, as indicated. The testes were stained with the Hoechst (red) and Phalloidin (green/ FIRE). F-actin intensities in the actin caps are depicted using the heat map in A'. Arrows and arrowheads indicate caplets and baskets, respectively. Yellow arrowheads mark unusual actin extensions from the basket. **C-D)** Low magnification image of the testis (C) and TE (D) from *PpY> Cdc 42 RNAi* background. The dashed yellow circle marks disrupted sperm heads.

**O-O')** *Rac1p-Rac-GFP* expression from the endogenous promoter in the HCC, along with Hoechst (blue) and Phalloidin (green). Note that Rac1-GFP localised surrounding the NB and overlapping with the basket domain (arrowhead)

**P-Q)** Box plots depict the number of intact NBs (P) and progressed ICs (Q) in different genetic backgrounds. The pairwise significance with respect to the control value was estimated using One-way Anova® and Mann-Whitney U tests, and the p-values- * <0.05, ** <0.01, *** < 0.001 are indicated on each box.

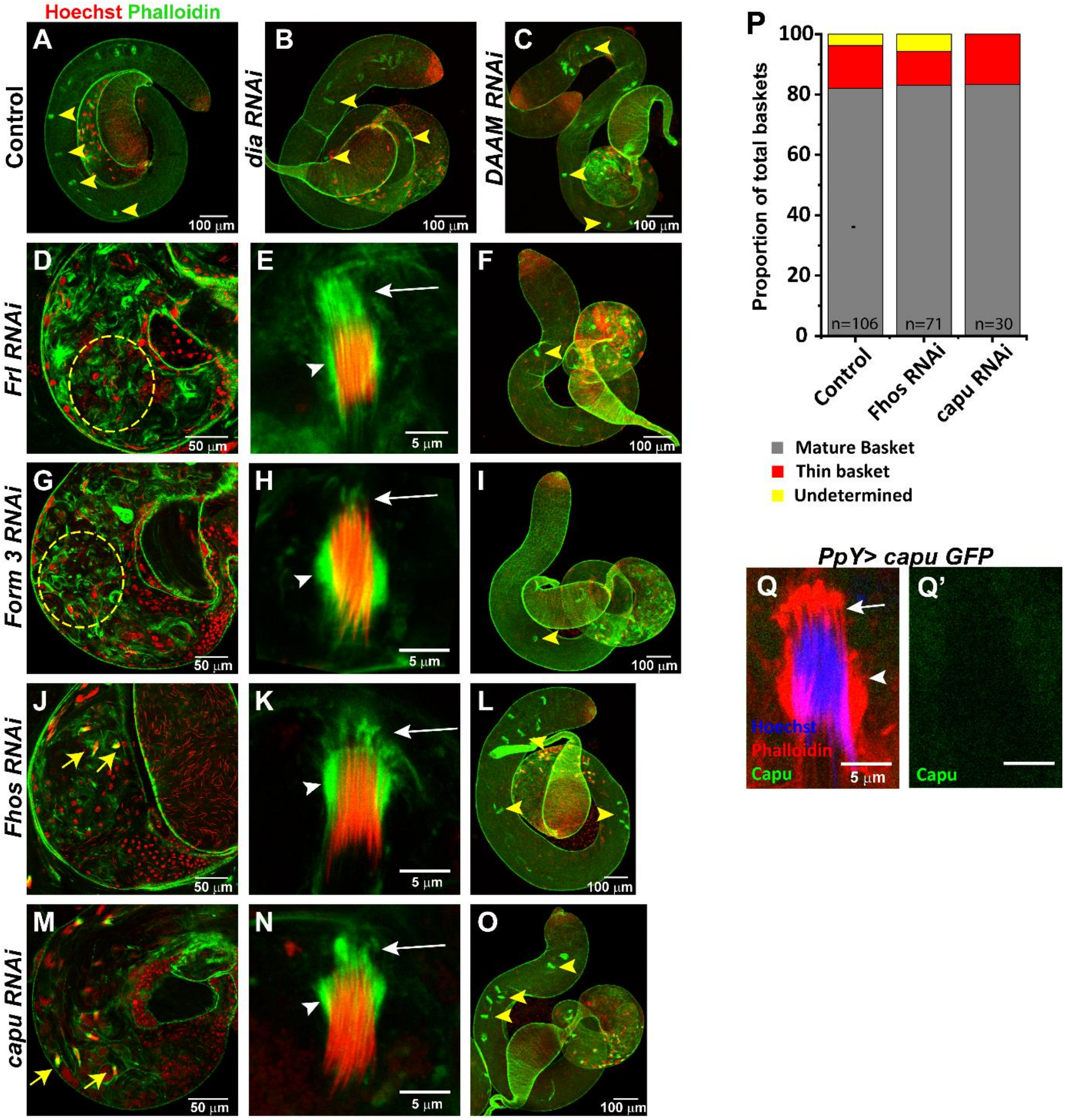


**Figure S5- Fhos and Capuccino are not involved in the basket assembly.**

**A-C)** Low magnification confocal images of entire testes of Control (A), *dia RNAi* (B), and *DAAM RNAi* (C) backgrounds, stained with Hoechst (red) and Phalloidin (green). Yellow arrowheads mark the progressed ICs.

**D-O)** Confocal images of *Frl* (D-F), *Form3* (G-I), *Fhos* (J-L) and *capu* (M-O) *RNAi* testes, depicting TE region (first column), intact NBs with associated actin caps (middle column) and entire testes representing progressed ICs (right-most column). Yellow arrows indicate intact NBs in TE; yellow dashed circle marks disrupted sperm heads; yellow arrowheads depict progressed ICs; white arrows mark caplets and white arrowheads mark basket.

**P)** Stacked bar plots indicates the relative distribution of mature baskets (grey) and thin baskets (red) in the *Fhos* and *capu RNAi* backgrounds. The number of NBs were too few in the *Frl* and *Form3 RNAi* backgrounds for similar quantification. Significance was calculated using Fisher's exact test. 'n' values are indicated on each bar.

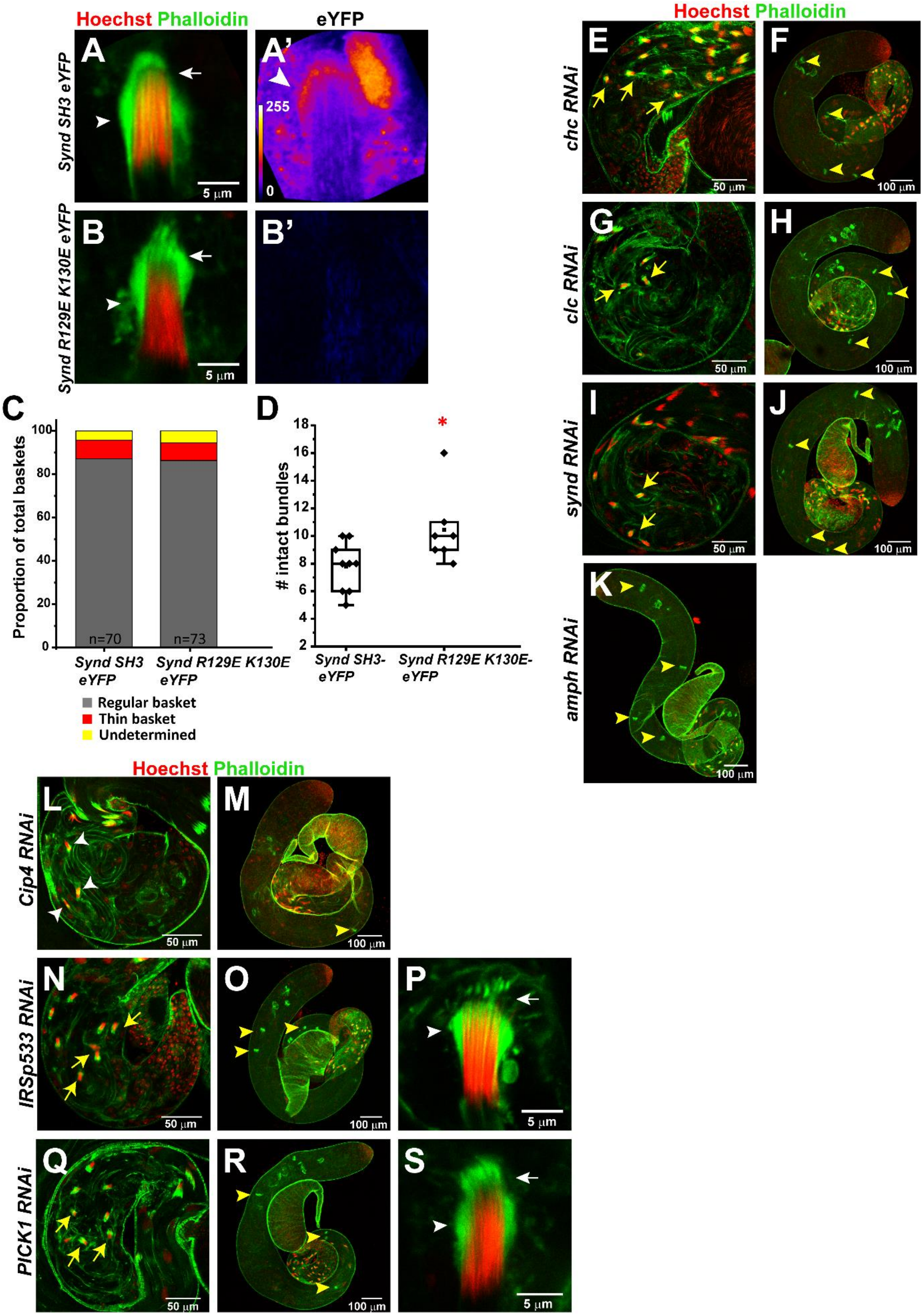
Hoechst Phalloidin
eYFP
A
A'
B
B'
Synd SH3 eYFP
Synd R129E K130E eYFP
255
0
5 µm
C
D
Proportion of total baskets
0
20
40
60
80
100
n=70
n=73
Synd SH3 eYFP
Synd R129E K130E eYFP
Regular basket
Thin basket
Undetermined
### intact bundles
2
4
6
8
10
12
14
16
18
*
Synd SH3-eYFP
Synd R129E K130E-eYFP
Hoechst Phalloidin
E
F
G
H
I
J
K
chc RNAi
clc RNAi
synd RNAi
amph RNAi
50 µm
100 µm
Hoechst Phalloidin
L
M
N
O
P
Q
R
S
Cip4 RNAi
IRSp533 RNAi
PICK1 RNAi
50 µm
100 µm
5 µm

**Figure S6- With the exception of Amph, loss of Synd and other BAR domain proteins does not affect spermatid bundling.**

**A-B')** NB from *PpY> Synd SH3-eYFP* (A-A') and *PpY> Synd R129E K130E-eYFP* (B-B') expressing testes, stained with Hoechst (red) and Phalloidin (green). Arrow and arrowheads mark caplets and basket, respectively. A' and B' represent the eYFP intensity (FIRE LUT) according to the heat map in A'. Arrowhead in A' marks the enrichment of *Synd SH3-eYFP* in the basket.

**C-D)** Stacked bar graph representing the basket phenotypes (C), and dot and box plots representing the number of intact NBs (D) in *Synd SH3-eYFP* and *Synd R129E K130E-eYFP* expressing testes. Significance was calculated using Fisher's exact test and Mann Whiteney U test, respectively. p-values- * <0.05 is indicated on the box.

**E-S)** Confocal images of the TE region (E, G, I, L, N, Q) and low magnification images of the entire testes (F, H, J, K, M, O, R) of *chc RNAi*, *clc RNAi*, *synd RNAi*, *amph RNAi*, *Cip 4 RNAi*, *IRSp53 RNAi and PICK1 RNAi* expressing testes. Yellow arrows mark intact NBs with associated actin caps, and yellow arrowheads mark progressed ICs. White arrowheads in L mark NBs with perturbed actin caps. P and S represent high magnification images of NBs from *IRSp53 RNAi and PICK1 RNAi* testes, respectively. Arrows and arrowheads mark baskets and caplets, respectively.

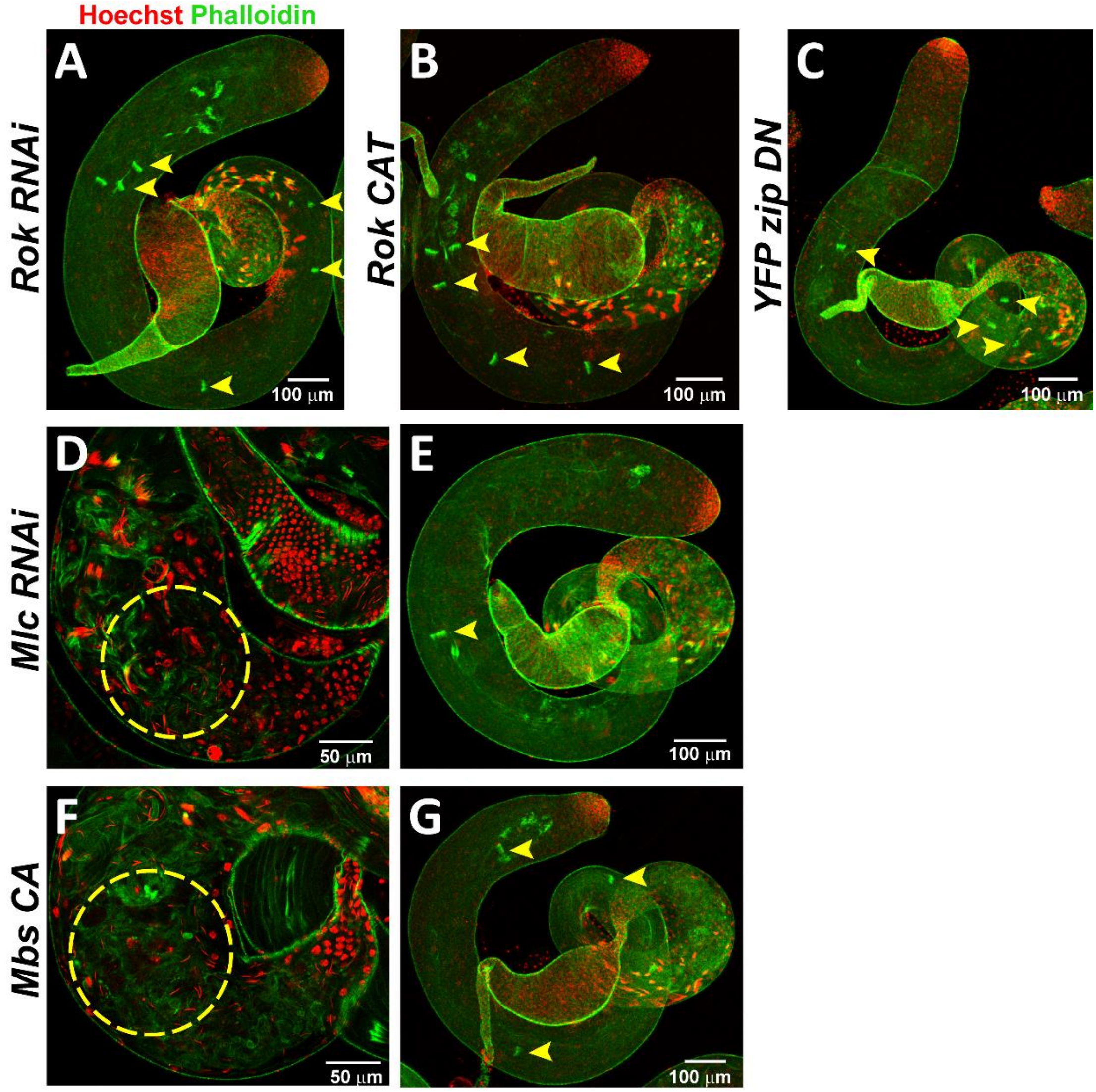


**Figure S7- Loss of bundled NBs and progressed ICs in *Mlc RNAi* and *Mbs CA* backgrounds**

**A-C)** Low magnification of entire testes of *Rok RNAi* (A), *Rok CAT* (B) and *YFP::zip DN* (C), stained with Hoechst (red) and Phalloidin (green), representing the progressed ICs (yellow arrowheads).

**D-G)** Confocal sections of the TE region (D, F), and low magnification image of entire testes (E, G) of *Mlc RNAi* (D-E) and *Mbs CA* (F-G), stained with Hoechst (red) and Phalloidin (green). Yellow dashed circles mark disrupted bundles and yellow arrowheads mark progressed ICs.

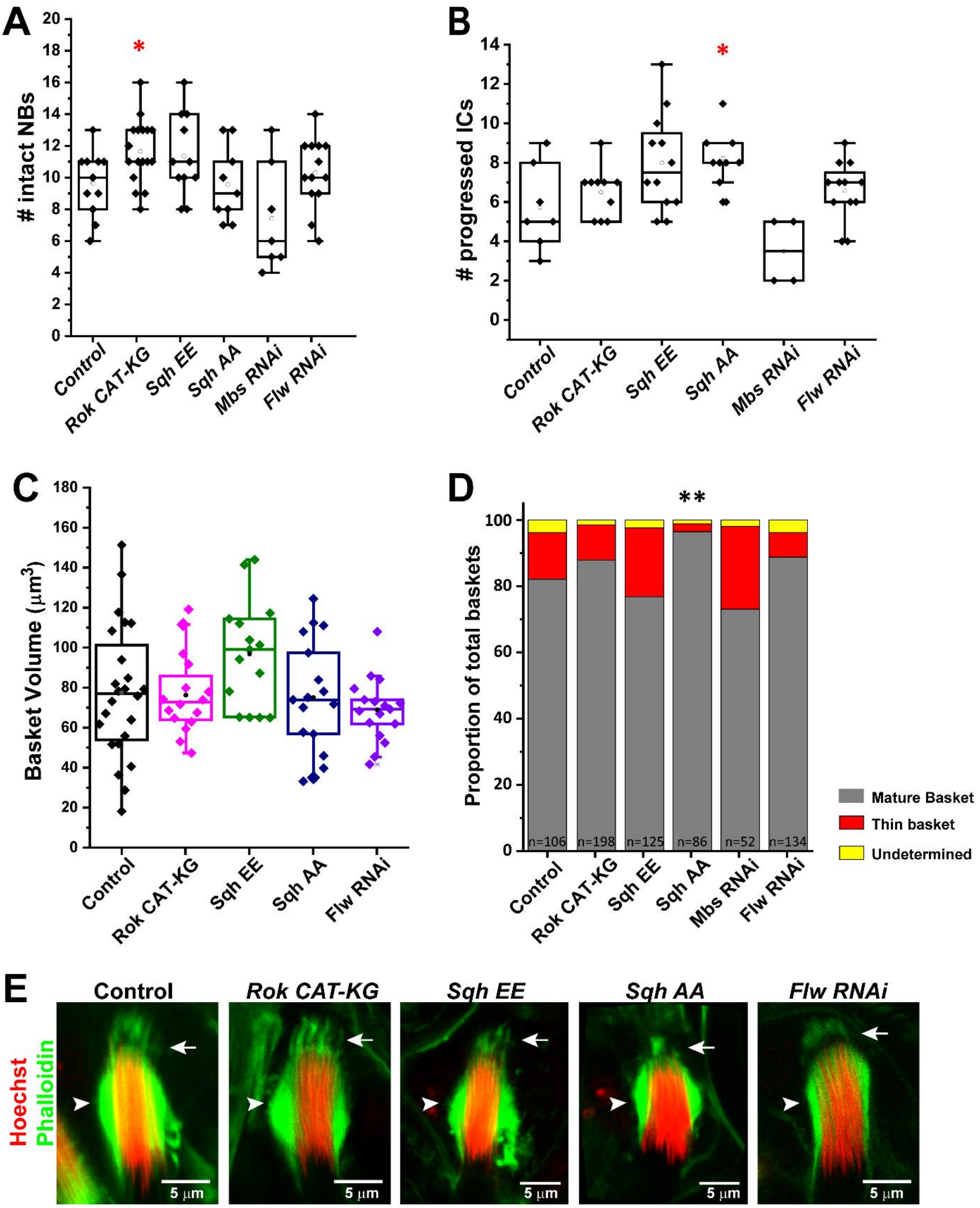


**Figure S8- Genetic perturbations to myosin activity, in the cyst cells, do not perturb the basket morphology.**

**A-C)** Box and dot plots representing the number of intact NBs (A), progressed ICs (B), and the volume of Phalloidin in the basket (C) in Control, *Rok CAT-KG*, *Sqh EE*, *Sqh AA*, and *Flw RNAi* expressing testes. One-way Anova® and Mann-Whitney U tests, respectively, and the p-values- * <0.05, ** <0.01, *** < 0.001 are indicated on each box.

**D)** Stacked bar column representing the frequency of occurrence of different basket phenotypes. The significance w.r.t control was calculated, using the Fisher's exact test, and the p-values- * $<0.05$, ** $<0.01$, *** $< 0.001$ are indicated on the column

**E)** Confocal images representing the actin cap phenotype in Control, *Rok CAT-KG*, *Sqh EE*, *Sqh AA*, and *Flw RNAi* expressing testes, stained with Hoechst (red) and phalloidin (green). Arrows and arrowheads mark caplets and baskets respectively.

**Movie legends**

**Movie S1-** 3D revolving movie depicting an NB (red) with associated actin cap (green) in WT testes. Rendered volumes of the entire actin cap is depicted in cyan, and basket domain is depicted in magenta. Stray F-actin detected in the ROI is depicted in dark blue. Such stray puncta were not included for the volume analysis.

**Movie S2-** Time lapse imaging of *sqh::utr-GFP; ProtB-dsRed* testes revealed that the basket (green) undergoes cycles of contraction and relaxation along the long axis of the NB (red). Double headed arrow tracks the length of the basket; white line marks the caudal end of the NB; arrowhead and arrow marks basket and caplet respectively. Scale- 5 µm.

**Movie S3-** Laser ablation of the basket leads to disruption of the actin cap (green) and the NB (red) in *sqh::utr-GFP; ProtB-dsRed* testes. The boxed area represents the ablated region in the post-ablation frame. An 800 nm IR laser was used at an intensity of 408.2 mW (5 iterations). Arrowhead and arrow marks basket and caplet respectively. Scale- 5 µm.

**Movie S4-** Control laser ablation in the cytoplasm of the HCC does not lead to a perturbation to the actin cap (green), or the NB (red). The boxed area represents the ablated region in the post-ablation frame. Arrowhead and arrow marks basket and caplet respectively. Scale- 5 µm.

**Movie S5-** FRAP (Fluorescence Recovery After Photobleaching) of the basket, in the boxed area, leads to partial recovery of the F-actin intensity. Arrowhead and arrow marks basket and caplet respectively. Scale- 5 µm.

**Movie S6-** Time-lapse imaging reveals that F-actin tubules (open arrowheads) periodically emanate from the basket in *sqh::utr-GFP; ProtB-dsRed* testis. Arrowhead and arrow marks basket and caplet respectively. Scale- 5 µm.

**Movie S7-** Control DMSO treatment of *sqh:sqh-GFP; ProtB-dsRed* testis. Sqh (green) localisation at the basket, and NB organisation red) is maintained throughout the period of imaging. Arrowhead marks basket. Scale- 5 µm.

**Movie S8-** Treatment of *sqh::sqh-GFP; ProtB-dsRed* testis with 200 µM Rockout. Over time, Sqh (green) localisation is lost from around the basket, and concomitantly the NB (red) organisation is perturbed. Arrowhead marks basket. Scale- 5 µm.

**Movie S9-** Control DMSO treatment of *sqh::utr-GFP; ProtB-dsRed* testis. Actin cap (green) and NB (red) are unperturbed by DMSO treatment. Arrowhead and arrow marks basket and caplet respectively. Scale- 5 µm.

**Movie S10-** *sqh::utr-GFP; ProtB-dsRed* testis treated with 200 µM Rockout. Rockout treatment leads to shearing of the basket and NB. Arrowhead and arrow marks basket and caplet respectively. Scale- 5 µm.

**Table S1-** List of *Drosophila* stocks used in the study

| Stock | Details |
|---|---|
| *CsBz* | |
| *SG18.1 Gal4* | BL-6405, BDSC |
| *PpY55A Gal4* | Dr. Eyal Schejter, Weizzmann Institute, Israel |
| *ProtB-dsRed* | Prof. John Belote, Syracuse University, USA |
| *Sqh::Utr-GFP* | Prof. Andrea Brand, The Gurdon Institute, UK |
| *UAS GFP-Utr* | Prof. Thomas Lecuit, Aix Marseille Université, CNRS, France |
| *UAS eGFP* | BL-5431, BDSC |
| *UAS Rho 1 RNAi* | GD-12734, VDRC |
| *UAS Rho 1 N19 DN* | BL-7328, , BDSC |
| *UAS Rho 1 V14 CA* | BL-8144, BDSC |
| *UAS Rho 1* | BL-28872, BDSC |
| *UAS Cdc 42 RNAi* | BL-29006, BDSC |
| *UAS Cdc 42 L89 DN* | BL-6289, BDSC |
| *UAS Rac 1 RNAi* | BL-28985, BDSC |
| *UAS Rac 1 N17 DN* | BL-6292, NCBS Fly Facility, Bengaluru, India |
| *UAS Rac 1 L89 DN* | BL-6290, BDSC |
| *UAS Rac 2 RNAi* | GD-28926, VDRC |
| *Rac 1p GFP Rac1* | BL-52284, BDSC |
| *UAS dia RNAi* | BL-28541, BDSC |
| *UAS DAAM RNAi* | BL-39058, BDSC |
| *UAS Frl RNAi* | BL-32447, BDSC |
| *UAS Form 3 RNAi* | BL-32398, BDSC |
| *UAS Fhos RNAi* | BL-31400, BDSC |
| *UAS capu RNAi* | BL-32922, BDSC |
| *UAS Capu GFP* | BL-24763, BDSC |
| *UAS eGFP Clc* | BL-7107, BDSC |
| *UAS Synd GFP* | Prof. Vimlesh Kumar, IISER Bhopal, India |
| *UAS chc RNAi* | Prof. Richa Ricky, IISER Pune, India |
| *UAS clc RNAi* | BL-27034, BDSC |

| | |
|---|---|
| *UAS Synd RNAi* | Prof. Richa Ricky, IISER Pune, India |
| *UAS Amph RNAi* | BL-53971, BDSC |
| *UAS Cip4 RNAi* | BL-31646, BDSC |
| *UAS IRSp53 RNAi* | GD-50019, VDRC |
| *UAS PICK 1 RNAi* | GD-22268, VDRC |
| *UAS Synd SH3 eYFP* | Prof. Richa Ricky, IISER Pune, India |
| *UAS Synd R129E K130E eYFP* | Prof. Richa Ricky, IISER Pune, India |
| *Sqh::Rok-GFP* | BL-52289, BDSC |
| *UAS zip-GFP* | Prof. Benny Shilo, Weizmann Institute, Israel |
| *sqh::sqh-GFP* | BL-57145, BDSC |
| *UAS Rok RNAi* | KK-104675, VDRC |
| *UAS Rok-CAT* | BL-6669, BDSC |
| *YFP-zip DN* | Prof. Andrea Brand, The Gurdon Institute, UK |
| *UAS Mlc RNAi* | GD-7916, VDRC |
| *UAS Mbs CA* | BL-63791, BDSC |
| *UAS Rok CAT-KG* | BL-6671, BDSC |
| *UAS Sqh EE* | BL-64411, BDSC |
| *UAS Sqh AA* | BL-64114, BDSC |
| *UAS Mbs RNAi* | BL-32516, BDSC |
| *UAS Flw RNAi* | BL-38336, BDSC |

BDSC- Bloomington *Drosophila* Stock Center, Indiana University, Indiana, USA

VDRC- Vienna *Drosophila* Resource Center, Vienna, Austria
